# Endogenous OptoRhoGEFs reveal biophysical principles of epithelial tissue furrowing

**DOI:** 10.1101/2024.05.12.593711

**Authors:** Andrew D. Countryman, Caroline A. Doherty, R. Marisol Herrera-Perez, Karen E. Kasza

## Abstract

During development, epithelia function as malleable substrates that undergo extensive remodeling to shape developing embryos. Optogenetic control of Rho signaling provides an avenue to investigate the mechanisms of epithelial morphogenesis, but transgenic optogenetic tools can be limited by variability in tool expression levels and deleterious effects of transgenic overexpression on development. Here, we use CRISPR/Cas9 to tag *Drosophila* RhoGEF2 and Cysts/Dp114RhoGEF with components of the iLID/SspB optogenetic heterodimer, permitting light-dependent control over endogenous protein activities. Using quantitative optogenetic perturbations, we uncover a dose-dependence of tissue furrow depth and bending behavior on RhoGEF recruitment, revealing mechanisms by which developing embryos can shape tissues into particular morphologies. We show that at the onset of gastrulation, furrows formed by cell lateral contraction are oriented and size-constrained by a stiff basal actomyosin layer. Our findings demonstrate the use of quantitative, 3D-patterned perturbations of cell contractility to precisely shape tissue structures and interrogate developmental mechanics.

## Introduction

Embryonic development is an inherently biophysical process, requiring the integrated action of protein regulators and mechanical effectors to physically sculpt an embryo into a mature animal. Therefore, a proper understanding of embryogenesis requires dissecting both the biochemical pathways that direct morphogenesis and the mechanical behavior of the cells and tissues that enact the developmental program (Collinet and Lecuit 2021). Across many animals, early development is defined by the sculpting of epithelia (Gilmour et al., 2017). Although epithelial tissues are effectively 2D sheets, precise subcellular patterning of active stresses and mechanical properties can shape them into complex 3D morphologies (Pilot and Lecuit 2005, Zartman and Shvartsman 2010, Tozluoǧlu and Mao 2020).

A well-studied mechanism to sculpt epithelial sheets is apical constriction, which directs tissue furrowing and invagination in a variety of contexts (Baker and Schroeder 1967, Martin et al., 2009, Eiraku et al., 2011). This process is characterized by a contractile apical actomyosin meshwork driving constriction of apical cell surfaces and expansion of basal cell surfaces (Martin and Goldstein 2014). However, recent theoretical and experimental work has additionally implicated lateral actomyosin contractility and cell apical-basal shortening in morphogenesis (Kondo and Hayashi 2015, Ioannou et al., 2020). Epithelial furrowing driven by lateral cortical actomyosin plays roles in *Drosophila* cephalic furrow formation (Spencer et al., 2015, Eritano et al., 2020) and wing disc morphogenesis (Sui et al., 2018), as well as ascidian endoderm invagination (Sherrard et al., 2010). More generally, lateral actomyosin contractility gives rise to a columnar-to-cuboidal transition during *Drosophila* pupal wing shaping (Athilingam et al., 2021) and apoptosis-coupled tissue bending during *Drosophila* leg disc folding (Monier et al., 2015). It is unknown why developing embryos employ different furrowing mechanisms in different contexts, or what functional consequences these may have. Additionally, while apical constriction provides a clear polarity-based cue guiding the direction of furrowing, it is unclear how lateral contraction translates cell shortening into a properly oriented furrow.

Although different morphogenetic processes are driven by distinct patterns of tissue stresses and mechanics, these are largely downstream of the widely conserved Rho signaling pathway (Etienne-Manneville and Hall 2002, Simões et al., 2006). In the Rho pathway, RhoGEFs and RhoGAPs cycle Rho GTPases between their active and inactive states to coordinate downstream actomyosin activity and direct the mechanical behavior of the epithelium. Different RhoGEFs and RhoGAPs are employed in a context-specific manner to initiate Rho signaling, but they converge on tuning the behavior of just a few classes of mechanically relevant molecules (Denk-Lobnig and Martin 2019). RhoGEF and RhoGAP activities must be balanced to carry out cell mechanical behaviors, and many are autoinhibited to prevent off-target activity (Mikawa et al., 2008, Müller et al., 2020). Additionally, some RhoGEFs are known to engage in positive feedback interactions with active Rho1 (Chen et al., 2010, Medina et al., 2013, Rich et al., 2020, Lin et al., 2022). Therefore, it is not clear to what extent tissues can modulate their shapes by controlling activity levels of a particular RhoGEF, or to what extent RhoGEFs act as on/off switches to enact tissue morphogenetic processes.

The *Drosophila melanogaster* embryo is a powerful model system for understanding how Rho signaling directs morphogenesis in a variety of contexts. RhoGEF2 and Cysts/Dp114RhoGEF, hereafter referred to as Cysts, are two RhoGEFs that play important roles in early fly development. During ventral furrow formation, RhoGEF2 directs apical actomyosin accumulation and apical constriction (Häcker and Perrimon 1998, Kölsch et al., 2007). During cephalic furrow formation, RhoGEF2 is a major driver and Cysts is a minor driver of cell shortening (Popkova et al., 2024). During germband extension, RhoGEF2 mediates apical contractility while Cysts mediates junctional contractility (Garcia De Las Bayonas et al., 2019, Silver et al., 2019). Thus, different RhoGEFs have overlapping and distinct roles in embryogenesis, and it remains unclear to what extent a single RhoGEF can direct a range of morphogenetic behaviors on its own.

Optogenetics has emerged as a valuable tool to precisely manipulate spatiotemporal patterns of Rho signaling and contractile stresses (Kennedy et al., 2010, Guntas et al., 2015). Optogenetic tools perturbing the Rho pathway have produced significant insight into embryogenesis by facilitating perturbation of developmental events and reconstitution of some morphogenetic processes (Izquierdo et al., 2018, Rich et al., 2020, Herrera-Perez et al., 2021). However, existing transgenic tools can be limited by variable expression levels, which hinder their quantitative use, as well as by artifacts of transgenic overexpression, which can unintentionally perturb development or compromise embryonic viability.

Here, we develop two new optogenetic tools based on the tagging of *Drosophila* RhoGEF2 and Cysts at their endogenous loci with a component of the light-activated iLID/SspB heterodimer, producing viable embryos and healthy flies. We present an activation quantification strategy and uncover a dose-dependent relationship between RhoGEF recruitment and epithelial furrowing and bending. We find that at the onset of gastrulation, furrow asymmetry and furrow size constraints are influenced by a stiff basal actomyosin network, and that disrupting this basal network via *scraps* RNAi alters these mechanical behaviors. Collectively, our results establish a framework for the precise optogenetic manipulation of tissue stresses and structures to interrogate biophysical mechanisms of morphogenesis.

## Results

### Development and characterization of two endogenous OptoRhoGEFs

We engineered an optogenetic system into *Drosophila melanogaster* based on the LOV2-derived iLID/SspB molecular switch, in which light exposure causes an iLID conformational change that promotes its heterodimerization with SspB (Guntas et al., 2015). We localize the iLID component to the cell membrane and attach the SspB component to a RhoGEF, so that upon blue light exposure, the RhoGEF is targeted to the membrane, where it locally activates the Rho signaling pathway and induces cellular contractility (Fig. 1A). In contrast to existing optogenetic tools targeting Rho signaling (Izquierdo et al., 2018, Rich et al., 2020, Herrera-Perez et al., 2021), here we used CRISPR/Cas9 to tag endogenous RhoGEF2 and Cysts loci with a SspB peptide, and optionally, a tagRFP-T sequence (Shaner et al., 2008), producing RhoGEF2-tgRFPt-SspB, RhoGEF2-SspB, SspB-tgRFPt-Cysts, and SspB-Cysts constructs (Gratz et al., 2014). The resulting flies were homozygous-viable. We then co-expressed these in the blastoderm with a transgenic plasma membrane-localized iLID construct - either Venus-iLID-CaaX, mCherry-iLID-CaaX, or iLID-CaaX, depending on the fluorescent structures being visualized. When an SspB-bound GEF and an iLID-CaaX construct are co-expressed, they form the complete optogenetic tool, which we refer to here as OptoGEF2 and OptoCysts.

**Figure 1.**
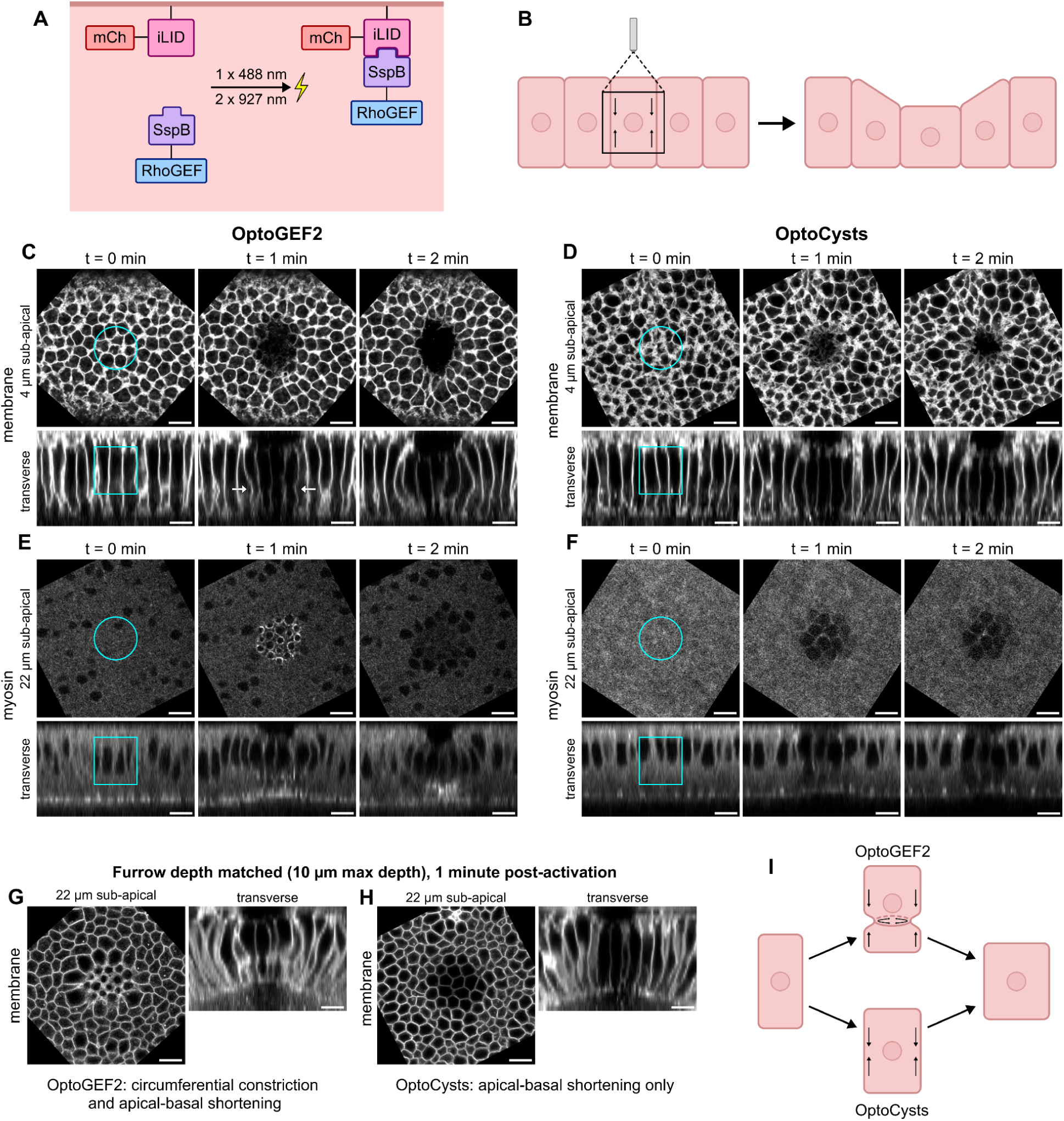
Lateral optogenetic recruitment of RhoGEF2 or Cysts results in tissue furrowing via cell apical-basal shortening. **(A)** Schematic: upon absorbing one 488-nm photon or two 927-nm photons, a membrane-bound iLID peptide undergoes a conformational change that promotes its binding to a GEF-bound SspB peptide, recruiting the GEF to the cell membrane. **(B)** Schematic: lateral RhoGEF recruitment is expected to induce apical-basal shortening and formation of a shallow furrow. **(C, D)** Two-photon lateral activation of embryos co-expressing transgenic mCherry-iLID-CaaX and 2 copies of endogenous RhoGEF2-tgRFPt-SspB (OptoGEF2, C) or transgenic mCherry-iLID-CaaX and 2 copies of endogenous SspB-tgRFPt-Cysts (OptoCysts, D) results in epithelial furrowing via lateral contraction. Arrows in (C) highlight in-plane constriction behavior. **(E)** Two-photon lateral activation of embryos co-expressing RhoGEF2-SspB (2 copies), Venus-iLID-CaaX, and sqh-mCherry (an mCherry-tagged *Drosophila* myosin II regulatory light chain) is associated with assembly of myosin II into transient ring-like structures at lateral interfaces. **(F)** Two-photon lateral activation of embryos co-expressing SspB-Cysts (2 copies), Venus-iLID-Caax, and sqh-mCherry is associated with localization of myosin II at polygonal cell outlines. **(G, H)** In depth-matched lateral activation experiments where cells shorten by 10 µm, transient circumferential constriction is observed in OptoGEF2 (G), but not OptoCysts (H), embryos (visualized with a mCh-iLID-CaaX membrane marker). **(I)** Schematic: Lateral activation results in persistent apical-basal shortening in both OptoGEF2 and OptoCysts embryos, but transient circumferential constriction only in OptoGEF2 embryos. For in-plane cross-sections, anterior is left and posterior is right. For transverse cross-sections, the apical tissue surface is oriented up and the basal surface is oriented down. Scale bars are 10 µm.

OptoGEF2 and OptoCysts embryos are viable, and their viability is not strongly affected by ambient light incubation or by being homozygous or heterozygous for the GEF construct (Fig. S1A). SspB-tagged RhoGEF2 and Cysts levels are highly consistent across embryos, displaying relative fluctuations of only 10-15%, in contrast to the UAS-driven iLID construct, which exhibits relative fluctuations of 45% (Fig. S1B, S1C). This makes these endogenously tagged RhoGEFs ideal for performing reproducible, quantitative optogenetics experiments.

One-photon (1P) activation of OptoGEF2 or OptoCysts with 488-nm light illumination, either globally or locally, led to GEF relocalization to the plasma membrane (Figs. 1A, S2A, S2B). However, to achieve subcellular recruitment, a two-photon (2P) activation protocol is required (Izquierdo et al., 2018, Krueger et al., 2019a). To overcome the inefficient 2P absorption of common optogenetic switches, such as iLID/SspB (Kinjo et al., 2019), we developed a strategy in which strong optogenetic recruitment can be driven by an inefficient activation process if the light-sensitive component of the optogenetic heterodimer is expressed in excess. To illustrate this general strategy, instead of examining our endogenously tagged RhoGEFs that are expressed at low levels, we visualized 2P recruitment in the more accessible, overexpressed OptoSOS system, also based on iLID/SspB heterodimerization (Johnson et al., 2017). OptoSOS uses a self-cleaving peptide to express the iLID and SspB components at roughly equimolar levels (Johnson et al., 2017). As LOV domains tend to show 800-950 nm 2P absorption (Homans et al., 2018), we tested these wavelengths. We find that impractically high 927-nm laser intensities are required to achieve visible OptoSOS 2P recruitment, resulting in heavy bleaching and eventual tissue ablation (Figs. S2C, S2E). However, when we co-expressed OptoSOS with a transgene encoding an additional copy of the light-sensitive iLID membrane anchor, we were able to observe clear membrane recruitment at modest 927-nm laser intensities (Fig. S2F). Based on this strategy, in OptoGEF2 and OptoCysts, we combine our endogenous SspB-tagged RhoGEF constructs with an overexpressed iLID anchor to provide the advantages of both endogenous protein manipulation and localized two-photon activation.

### Lateral RhoGEF recruitment causes tissue furrowing via cell apical-basal shortening

Based on previous work employing a transgenic optogenetic tool using the catalytic DH/PH domains of RhoGEF2, we hypothesized that 2P activation of a lateral cell volume would recruit endogenous RhoGEF2 or Cysts to lateral cell interfaces, driving cell apical-basal shortening and producing a shallow furrow (Fig. 1B) (Sui and Dahmann 2020, Popkova et al., 2024). Here, we define “shortening” as a reduction in the distance between the apical and basal cell surfaces, and “furrowing” as an overall movement of cells toward the basal surface of the epithelium, producing a pocket on the apical surface. This is distinct from invagination, which additionally requires cells to be internalized. We note that gastrulation in *Drosophila* and other animals requires furrowing and invagination toward the embryo interior to properly separate germ layers- namely, the ectoderm from the mesoderm and endoderm.

We found that 927-nm lateral activation in embryos finishing cellularization resulted in cell shortening and tissue furrowing in both OptoGEF2 and OptoCysts embryos, with a stronger response in OptoGEF2 (Fig. 1C, 1D). Shortening was also induced by 800, 850, or 900-nm activation (Fig. S3). Thus, despite their frequent autoinhibition (Müller et al., 2020), manipulating the spatial localization of endogenous RhoGEFs can drive ectopic tissue shape changes. Moreover, creating RhoGEF membrane anchoring sites is sufficient to induce a contractile response, and no active relocalization mechanisms are required.

Although activation of either OptoGEF2 or OptoCysts induced persistent basal-directed cell shortening, the two tools displayed different transient cell deformation states. At 1 minute post-activation, OptoCysts embryos showed smooth lateral interfaces, while OptoGEF2 embryos showed increased lateral curvature and unexpected circumferential constriction at a sub-nuclear plane, ∼22µm below the apical surface (Figs. 1C and 1D, lower middle panels). This temporarily produced cells with an hourglass-like lateral contour only in OptoGEF2 embryos (Figs. 1C and 1D, lower middle panels). Comparing tissue behaviors in OptoGEF2 and OptoCysts embryos that both form a 10-µm deep furrow, circumferential constriction is only seen in OptoGEF2, suggesting that the phenotype is not due to the strength of the mechanical response (Figs. 1G, 1H). Imaging nonmuscle myosin II (hereafter referred to as myosin), we found that Cysts recruitment resulted in accumulation of myosin at lateral cell-cell interfaces, as expected, while RhoGEF2 recruitment additionally organized myosin into ring-like structures at the basal ends of nuclei (Figs. 1E and 1F, upper middle panels). At two minutes post-activation, OptoCysts myosin recruitment is limited to lateral interfaces, while OptoGEF2 embryos have a distinct additional accumulation of myosin near the basal cell surface. (Figs 1E and 1F, lower right panels).

Given that RhoGEF2 localizes to microtubules (MTs) (Rogers et al., 2004) and that *Drosophila* blastoderm cells have prominent perinuclear MT bundles (Warn and Warn 1986, Callaini and Anselmi 1988), we imaged endogenous RhoGEF2 tagged with tdTomato to see if RhoGEF2 was concentrated around nuclei (Fig. S4). We found that RhoGEF2 displayed no overall enrichment at the level of nuclei (Fig. S4A), but it accumulated at membrane-adjacent puncta in perinuclear regions (Fig. S4B). It’s possible that RhoGEF2 is locally captured from these membrane-apposed puncta to drive the subnuclear constriction behavior.

To investigate the use of our endogenous optogenetic tools in alternative furrowing mechanisms, we also attempted to reconstitute apical constriction-based furrowing by selectively illuminating apical cell surfaces, as has been done with transgenic tools based on GEF catalytic domains (Izquierdo et al., 2018, Rich et al., 2020) (Fig. S5A). In OptoGEF2 embryos, we found that apical activation caused localized accumulation of myosin at apical caps (Fig. S5B), resulting in weak apical constriction and formation of a shallow furrow at the apical surface (Fig. S5C). However, this was accompanied by cell shortening and lateral expansion rather than basal expansion, inconsistent with traditional pictures of apical constriction-driven furrowing. This result is, however, similar to that observed after apical RhoGEF2(DHPH)-Cry2 recruitment in the *Drosophila* wing disc epithelium (Sui and Dahmann 2020). These results suggest that OptoGEF2 may have limitations associated with endogenous RhoGEF2 localization, full-length RhoGEF2 interacting partners, or RhoGEF2 autoinhibition.

Overall, lateral recruitment of either RhoGEF2 or Cysts causes cell shortening and tissue furrowing via lateral actomyosin accumulation. However, only RhoGEF2 recruitment induces transient circumferential constriction (Fig. 1I), possibly due to capture from nearby MTs.

### RhoGEF-driven cell shortening is dose-dependent

To dissect the mechanisms that specify the depth of an epithelial furrow during development, we asked whether we could precisely tune tissue furrowing by modulating the extent of OptoGEF2 or OptoCysts activation. To do this, we developed a pipeline to specify an input optogenetic dose and measure the resulting tissue furrowing response. After acquiring images of the unperturbed tissue (Fig. 2A), we scanned an IR laser (0-75 mW) for 40 seconds over a cylindrical lateral cell volume, 18 µm in diameter and extending from 5-25 µm below the apical cell surface (Fig. 2A’), activating the iLID in a 2P process. To avoid mechanical interference from endogenous ventral furrow formation, we activated the dorsal side of the embryonic primary epithelium. We activated cells when they were between 32-35 µm long, concomitant with the initiation of ventral furrow formation and with the transition from cellularization to gastrulation. We observe the tissue response for ∼5 minutes (Fig. 2A’’), up until the dorsal tissue starts to be compressed by the invaginating posterior midgut. We then segment the apical and basal surfaces of the tissue to extract relevant quantitative metrics of the furrowing response (Fig. 2A’’’).

**Figure 2.**
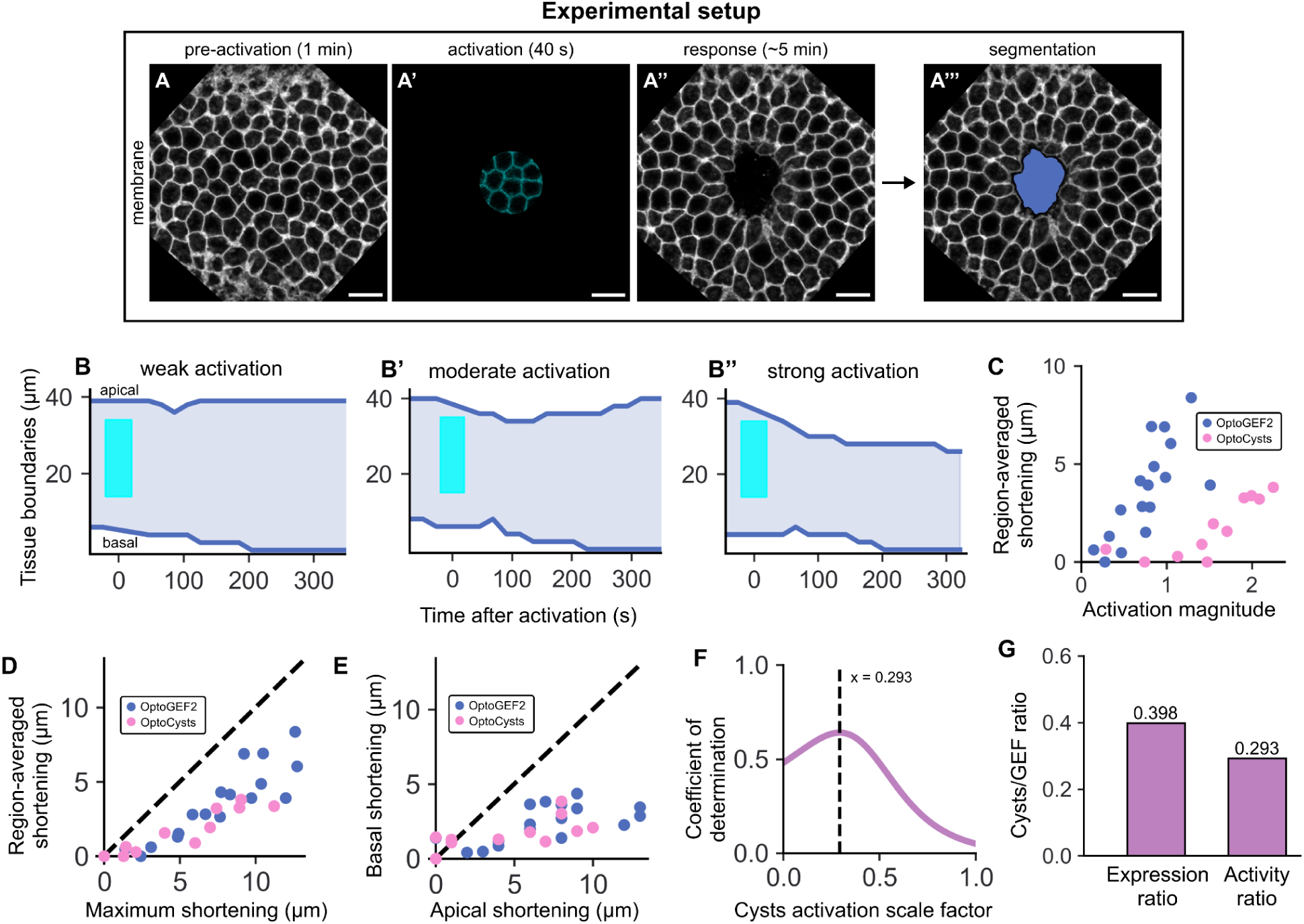
RhoGEF-driven cell shortening is dose-dependent. **(A)** Representative stills from each experimental step. After acquiring images of the unperturbed tissue (A), a 927-nm laser is scanned across a cylindrical cell volume measuring 18-µm in diameter and extending 5-25 µm beneath the apical tissue surface (A’). Tissue response to activation is recorded (A’’) and quantified via segmentation of the induced furrow (A’’’). In all steps, mCh-iLID-CaaX is the fluorescent membrane marker. **(B, B’, B’’)** Representative traces of OptoGEF2 embryo responses to different amounts of iLID activation. Upper curve represents the average position of the apical tissue surface within the activation region; bottom curve represents the average position of the basal surface within the activation region. Cyan region represents the 40-s activation step. **(B)** Weak activation protocols result in shallow furrows that quickly recover. **(B’)** Moderate activation protocols result in deeper, more persistent furrows. **(B’’)** Strong activation protocols result in deep furrows that do not recover by the end of the observation period (onset of gastrulation movements). Activation magnitudes are 0.15 (B), 0.46 (B’), 1.29 (B’’). **(C)** Using the emitted fluorescence of the mCherry-iLID-CaaX fusion construct during 927-nm activation (shown in A’) as a measure of activation magnitude, both OptoGEF2 and OptoCysts embryos show a dose-dependent response of region-averaged cell shortening to optogenetic stimulation. **(D)** The maximum cell shortening at any point in the activation region correlates with, but is larger than, the region-averaged shortening. **(E)** Shortening response is dominated by movement of the tissue apical surface. **(F)** Residuals of the fit to the combined OptoGEF2/OptoCysts shortening dataset are minimized when Cysts activation magnitudes are scaled by 0.29. **(G)** The difference in the activity of the OptoGEF2 and OptoCysts tools can largely be accounted for by differences in endogenous RhoGEF2/Cysts expression levels (measured in Fig. S1). Scale bars are 10 µm.

Based on the low variability in RhoGEF2 and Cysts expression levels (Fig S1C), we reasoned that the strength of the tissue response should be determined by the optogenetic dose we deliver - that is, by the number of iLID molecules that we photoconvert. This depends on two factors - the iLID expression level, which sets the base amount of inactive iLID molecules, and the photon flux passing through the illuminated cell volume, which depends on the activating laser power and determines what proportion of these iLID molecules are activated. For a 2P activation process, we expect the activation dose to scale quadratically with the activating laser power and linearly with the iLID expression level. Both of these factors are captured quantitatively in the fluorescence emitted by the mCh-iLID construct during our 40-second activation step, as mCherry is weakly excited by 927-nm light in a 2-photon process (Fig. 2A’). We thus use the relative iLID fluorescence from this activation step as a measure of activation magnitude, allowing us to compare tissue responses to different amounts of activation in different experiments and across embryos. This relies on the assumption that we are not saturating iLID activation, which holds here (Fig. S2).

First, we examine the average positions of the apical and basal tissue surfaces within the tracked activation region over time (Figures 2B-2B’’). Note that cells continue elongating by several microns until cellularization is fully completed, so that the final position of the basal surface is deeper than the initial position. Cellularization rates were the same between control, OptoGEF2, and OptoCysts embryos (Fig. S1D), and are all consistent with wild-type cellularization rates (Lecuit and Wieschaus 2000), indicating that differential cellularization rates do not play a role in tissue shape changes. We see that weak activation causes shallow, transient cell shortening (Fig. 2B), moderate activation elicits stronger, longer-lasting shortening (Fig 2B’), and strong activation produces deep furrows that persist through the end of the observation period (2B’’). This suggests that at sufficiently strong levels of activation, cell shortening can be converted into stable tissue furrowing.

Then, for each experiment, we extract the maximum amount of cell shortening, calculated as the largest difference between instantaneous cell height and reference cell height based on embryo-specific cellularization rates. Applying this analysis, we find a striking grading of cell shortening response across a broad range of optogenetic doses for both OptoGEF2 and OptoCysts (Fig. 2C). We note that our OptoCysts lines expressed the iLID component at higher levels than our OptoGEF2 lines, allowing us to sample higher activation magnitudes with the Cysts tool. RhoGEF2 dose-dependence has been described in migrating *Drosophila* PGCs (Lin et al., 2022), but to our knowledge has not been observed in the context of tissue morphogenesis. This dose-dependence suggests that although positive feedback likely amplifies RhoGEF activity in this context, it does not saturate it or cause switch-like behavior.

The shortening response was not uniform over the whole activation region, resulting in the maximum shortening within the region being slightly greater than the average shortening (Fig. 2D). Consistent with our qualitative observations in Figs. 1C and 1D, we found that shortening from the apical surface dominated the response, with shortening from the basal surface playing only a minor role (Fig. 2E). Using our dose-response curves, we calculated an “activity ratio” of OptoCysts compared to OptoGEF2 by finding the scaling factor which, when applied to the OptoCysts activation magnitudes, minimized the residuals to the joint OptoGEF2-OptoCysts shortening vs. activation magnitude linear fit (Fig. 2F). This activity ratio represents the relative dose-response sensitivity of the two tools and depends on the expression levels of the two RhoGEFs, their relative catalytic activities, and any potentially differential steric hindrance imposed by the optogenetic tags. We calculated an activity ratio of 0.29 (Fig. 2G), which is comparable to our measured Cysts/RhoGEF2 expression ratio of 0.40 (Fig. S1B), and to a Cysts/RhoGEF2 abundance ratio of 0.37 extracted from an embryo proteomics dataset (Casas-Vila et al., 2017), suggesting that differences in tool activity can largely be explained by differences in expression levels of the two GEFs.

Our dose quantification approach is validated by three additional observations. First, at constant activating laser power, shortening scales with iLID expression (Fig. S6A). Second, shortening behavior is much better accounted for by our activation magnitude parameter, which combines information about activating laser power and expression, than by activating laser power alone (Figs S6B, S6C). Finally, when we calculate our dose using an alternative measure, defined as the laser power squared times the relative iLID expression level, we also find dose-dependent shortening behavior (Fig S6D) and linear scaling with our activation magnitude parameter (Fig S6E). In alternative experimental setups where iLID fluorescence during activation can’t be measured, this alternative approach may prove useful.

Overall, our endogenous optogenetic tools allowed us to uncover a dose-dependent relationship between RhoGEF recruitment and furrow depth. This suggests that during development, furrow morphologies can be specified by modulating the amount of RhoGEF membrane anchoring sites. This is true of both RhoGEF2 and Cysts, suggesting that tunable furrowing is a common property of RhoGEFs, and that these two RhoGEFs overlap at least partially in the tissue deformations they are able to drive.

### Activation dose modulates the amount of RhoGEF recruited to the membrane

Although we found that furrow depth scales with our administered optogenetic dose, it is not obvious how this dose translates to actual changes in RhoGEF activity or localization. An unintuitive feature of a non-equimolar optogenetic switch in which the light-sensitive component is in excess (here, iLID) is that the timescale of SspB unbinding becomes decoupled from the timescale of iLID dark-state reversion (Natwick and Collins 2021). This is apparent via both an equilibrium argument, in which additional activated iLID beyond the amount required for near-full SspB translocation acts to increase the duration of full SspB recruitment, as well as a kinetic argument, in which excess unbound iLID at the membrane can trap SspB after it unbinds from an iLID molecule but before it diffuses into the cytoplasm (Natwick and Collins 2021). In OptoGEF2 and OptoCysts embryos, total iLID levels are in excess of RhoGEF levels, as previously described. However, given the demonstrated inefficiency of the 2P activation process (Kinjo et al., 2019) (Fig. S2), it’s uncertain if activated iLID is also in excess. Thus, depending on whether the RhoGEF or the active iLID is in excess, an increase in optogenetic dose will lead to either more RhoGEF molecules being recruited, or a roughly constant number of RhoGEF molecules being recruited for an increased duration (Fig. 3A).

**Figure 3.**
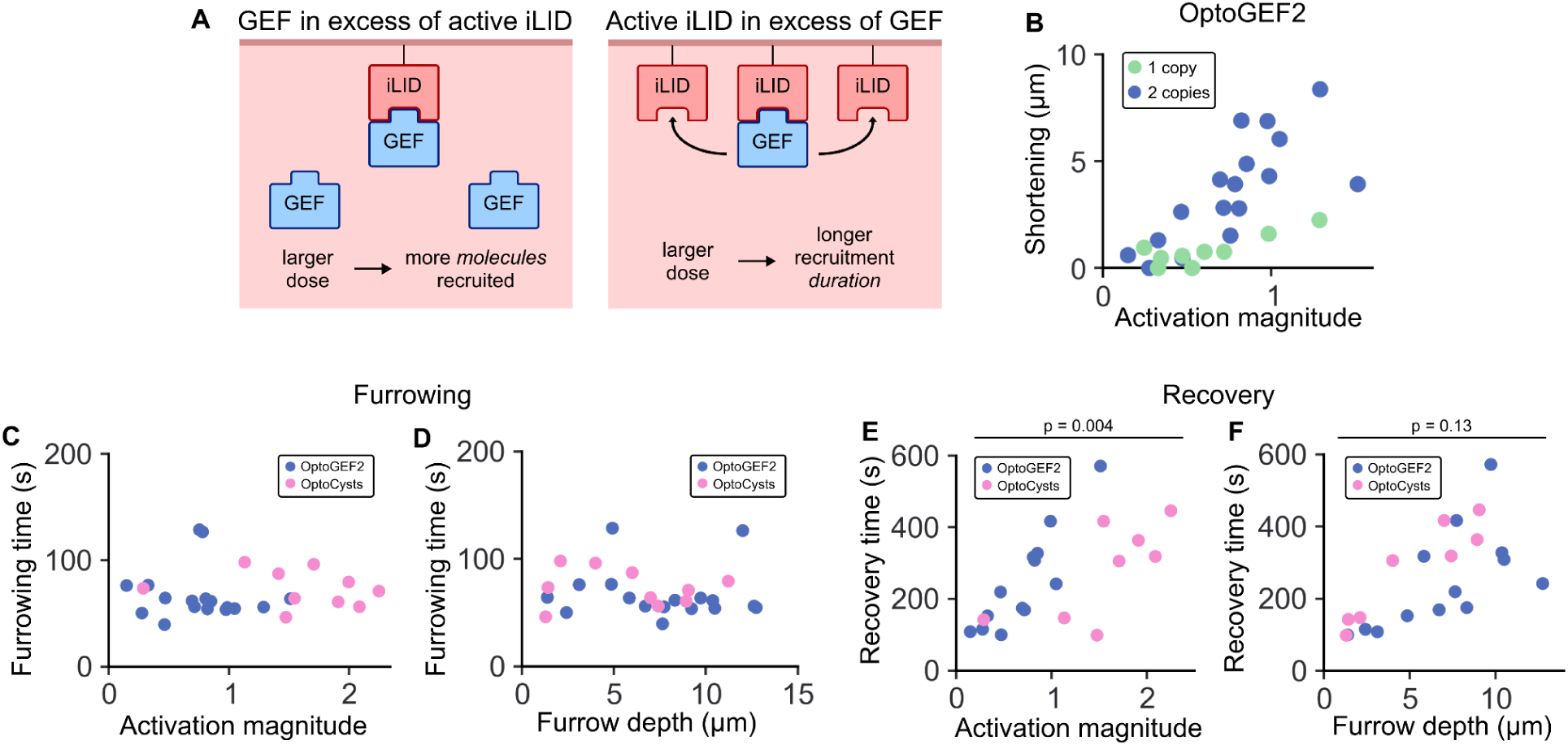
Activation dose modulates shortening via the amount of RhoGEF recruited to the membrane. **(A)** Schematic: depending on the relative abundance of iLID and SspB-bound RhoGEF, an increase in the number of activated iLID molecules can result in either increased magnitude or increased duration of GEF recruitment. **(B)** Embryos laid by mothers expressing mCh-iLID-CaaX and heterozygous for RhoGEF2-tgRFPt-SspB show a weaker shortening response than embryos laid by homozygous mothers. **(C, D)** The timescale of tissue furrowing is 70 seconds and is independent of either activation strength (C) or final furrow depth (D). **(E, F)** In experiments where furrows do recover, the timescale of recovery depends on the depth of the furrow (F) rather than the number of activated iLID molecules (E). P-values in E, F are calculated using a two-tailed t-test against the null hypothesis that the relationship between recovery time and activation magnitude (E) or recovery time and furrow depth (F) is the same for OptoGEF2 and OptoCysts embryos.

To dissect this, we measured the timescales of tissue furrowing and tissue recovery, the latter of which we could only quantify in furrows that relax. Surprisingly, we found that furrowing time was independent of the GEF activated, the activation magnitude, or the ultimate furrow depth, taking an average of 70 seconds across all experiments (Figs. 3C, 3D). The recovery time, on the other hand, did scale with activation magnitude and with furrow depth (Figs. 3E, 3F). Notably, the OptoGEF2 and OptoCysts recovery times only collapsed onto a single curve when plotted versus furrow depth, suggesting that the timescale of recovery is better explained by the final mechanical state of the furrow than by iLID reversion kinetics.

We argue that the optogenetic dose modulates the amount of RhoGEF at the membrane based on the following observations. First, if the mechanical behavior of the tissue is elastic, with a given amount of membrane-bound GEF corresponding to a given cell height strain, then we would only observe the dose-dependent furrow depth seen in Fig. 2C if our dose modulates the amount of GEF. Second, if the behavior of the tissue is viscous, with a given amount of membrane-bound GEF driving a given cell height strain rate, then we would only observe the constant furrow time seen in Fig. 3C if our dose changed the magnitude, rather than the duration, of GEF recruitment. Thus, in either case, our quantitative measurements are more consistent with the dose modulating the amount of membrane-bound GEF. This interpretation is supported by the furrow recovery timescales in Fig. 3F being mechanically determined and is consistent with, though not proven by, the observation that embryos heterozygous for RhoGEF2-tgRFPt-SspB show a weaker dose-dependent shortening response than their homozygous counterparts (Fig 3B). Although this is the most parsimonious interpretation, we cannot rule out potential complexities arising from biochemical feedback or endogenous protein localization patterns in determining the timescale or strength of the response. In summary, our optogenetic dose tunes furrow depth by changing the amount of RhoGEF2 recruited to the cell membrane, and furrow recovery dynamics scale with the depth of the induced furrow.

### Large regions of lateral Rho activation display disrupted furrowing and are associated with tissue bending

Comparing the wide, apical-constriction driven ventral furrow to the narrow, lateral contraction-driven cephalic furrow, we wondered how lateral contraction scales as a furrowing mechanism in the primary epithelium. To explore this, we increased activation region diameters up to 50 µm. Strikingly, we found that while 18-µm regions underwent coherent shortening toward the basal surface, eventually forming a furrow (Fig. 4A), 50-µm regions instead produced highly curved, concave tissues that predominantly shortened toward the apical surface and failed to form a productive furrow (Fig. 4C). This phenotype was also observed in OptoCysts embryos (Fig. S7A), although bending was not as strong. At intermediate region sizes, some tissues showed strong bending, but were still able to undergo net basal-oriented shortening to produce a furrow (Fig. 4B). Bent tissues showed a unique behavior in apical slices, with a central tissue plug surrounded by acellular perivitelline space (Fig. 4D). We used this behavior to classify embryos as either bending or non-bending. However, we did observe a continuous range of tissue morphologies from flat to curved, so bending behavior was not binary (Fig. 4E). Plotting bending phenotype as a function of activation region size and activation magnitude, we found that larger activation regions required smaller amounts of activation to exhibit bending (Fig. 4F).

**Figure 4.**
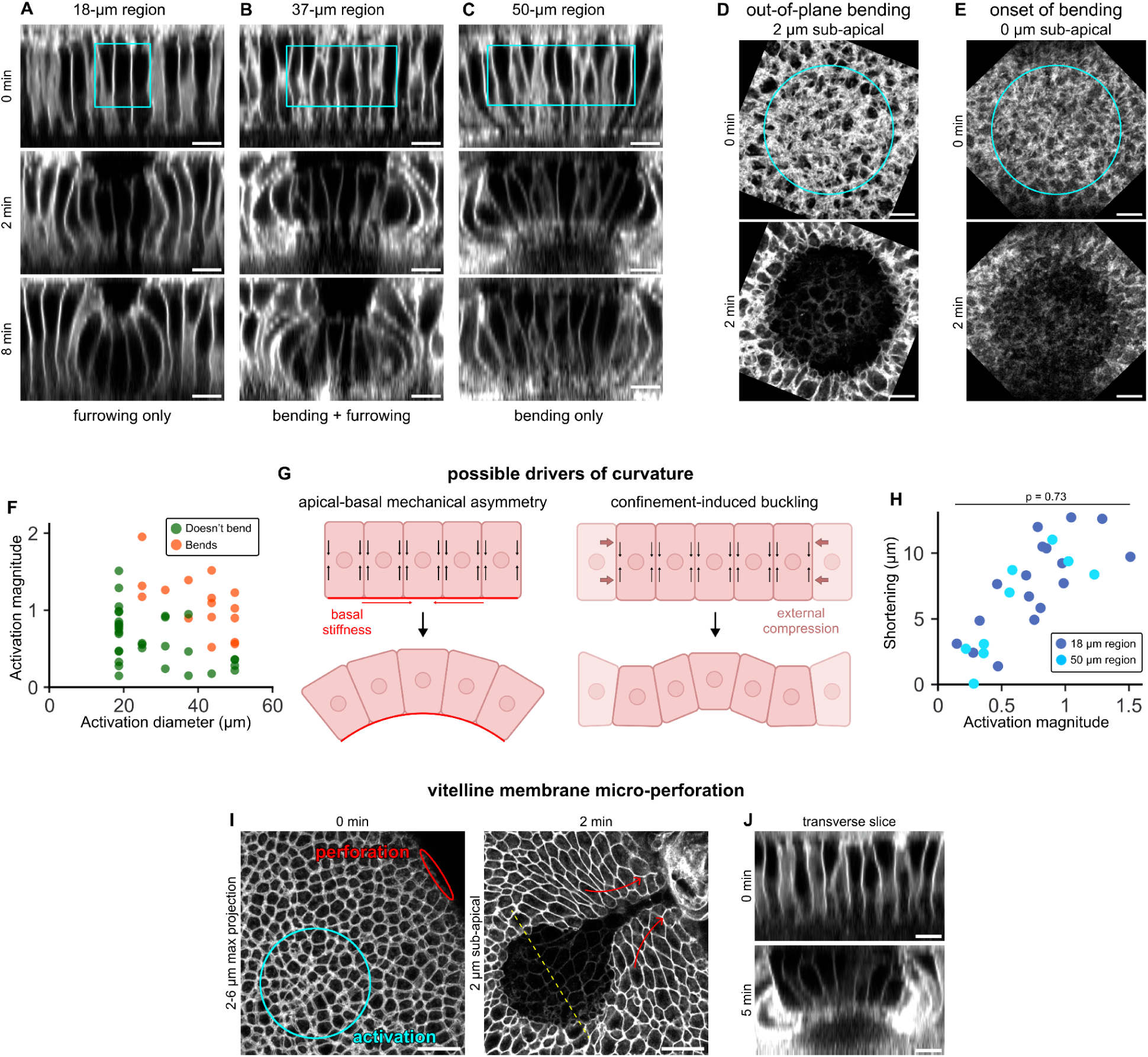
Large regions of lateral activation produce curved tissues that fail to form a furrow. **(A)** Activation of an 18-µm region results in tissue furrowing only. **(B)** Activation of a 37-µm region results in tissue furrowing and bending. **(C)** Activation of a 50-µm region results in tissue bending only. Activation magnitudes are 1.29 (A), 1.39 (B), 1.02 (C). **(D)** In-plane view of the stack shown in C, middle panel (50 µm region, activation magnitude = 1.02). Curved tissues are characterized by out-of-plane bending, which appears in an in-plane optical section as a central tissue plug surrounded by empty perivitelline space. **(E)** Apical slice of an activated tissue (50 µm diameter, activation magnitude = 0.36). Tissues adopt a continuous range of shapes between flat and curved. **(F)** Tissue bending depends on both the size of the activation region and the strength of activation, with larger regions showing curvature at smaller activation magnitudes. Bending is classified using the behavior described in (D). **(G)** Tissue curvature can arise from lateral contraction via either cell-intrinsic basal stiffness (left) or cell-extrinsic compressive forces (right). **(H)** Small and large activation regions result in the same maximum cell shortening, despite showing different tissue-scale furrowing and bending phenotypes (p = 0.73, two-tailed t-test against the null hypothesis that the relationship between cell shortening and activation magnitude is the same for 18-µm and 50-µm activation regions). **(I, J)** An 840-nm laser was used to activate a 52-µm region and then perforate a neighboring portion of the vitelline membrane, relieving hydrostatic pressure and tissue compression via outward-driven tissue flow. The curved shape of the activated region persists, as seen in in-plane (I) and transverse (J) slices. In all images, mCh-iLID-CaaX is the membrane marker. Scale bars in A-E, J are 10 µm. Scale bars in I are 20 µm.

We considered two mechanisms by which lateral contractility could produce curved tissues (Fig. 4G), both relying on volume conservation-based coupling between cell shortening and cell area expansion (Gelbart et al., 2012). In one model, tissue-intrinsic basal stiffness induces bending by restricting basal area expansion, producing wedge-shaped cells and net tissue curvature (Guerra Santillán et al., 2024a). In another model, tissue area expansion pushes against surrounding non-activated cells and the resulting external compressive force bends the activated tissue via a buckling or wrinkling instability (Hannezo et al., 2014, Andrenšek et al., 2023). The former model predicts curvature from tissue-intrinsic forces, while the latter model predicts curvature from tissue-extrinsic forces.

To distinguish between these potential mechanisms, we first note that curvature is not the result of a wrinkling instability, as activation regions measuring from 25-75 µm in diameter always resulted in only a single tissue crest, indicating that the bending pattern was not spatially periodic (Figs. 4F and S7B). Moreover, cell heights were equal in tissue grooves and crests, inconsistent with the groove-crest symmetry-breaking predicted during a wrinkling instability (Figs. S7C and S7D) (Andrenšek et al., 2023). In a buckling-type process, we expect a buildup of in-plane compressive stress during cell shortening that abruptly transitions to out-of-plane curvature once bending becomes energetically favorable. Here, we found that the scaling of maximum cell shortening with activation magnitude was equivalent in 18-µm and 50-µm regions (Fig. 4H) and that tissues displayed a continuous range of bending behaviors (Fig. 4E), both of which are inconsistent with a simple buckling process. Finally, relieving compressive stress and hydrostatic pressure via IR laser vitelline membrane micro-perforation (Popkova et al., 2024) does not relax tissue curvature, further suggesting that bending is a tissue-intrinsic process (Figs. 4I, 4J).

Overall, our observations rule out wrinkling and buckling as drivers of tissue curvature. This suggests that basal tissue stiffness is likely the source of tissue bending in our experiments. Here, bending obstructs productive furrowing in a size-dependent manner because large activation regions cause tissues to form an arc that subtends a larger angle, deviating more from an approximately flat morphology. Forcing the activated tissue to adopt this larger arc shape while still maintaining continuity with non-activated tissue along its peripheral basal surface causes the center of the activated region to bulge toward the embryo exterior instead of furrowing toward the embryo interior. We found that this bending behavior disrupts the ability of large regions of lateral contraction to form a coherent furrow, imposing an effective size constraint on furrows formed by this mechanism.

### Perturbing the basal tissue surface via *scraps/anillin* RNAi alters tissue furrowing and bending behaviors

During these studies, we identified multiple tissue behaviors that seemed consistent with the presence of a mechanically stiff basal layer in the primary epithelium. These included preferential cell shortening toward the basal surface for small activation regions (Fig. 2E) and tissue bending for large activation regions (Fig. 4C). Given that *Drosophila* cellularization is associated with a stiff, actomyosin-rich basal front (Warn and Magrath 1983, Warn and Robert-Nicoud 1990, Cheikh et al., 2023), we hypothesized that mechanical perturbation of the basal tissue surface would alter activation-induced tissue shape changes.

To test this, we repeated these experiments in a *scraps/anillin (scra)* RNAi background, in which cellularization is perturbed, characterized by destabilized yolk stalks and improper actomyosin organization at basal furrow canals (Thomas and Wieschaus 2004, Field et al., 2005, Goldner et al. 2023 *Preprint*). By the onset of gastrulation, rather than forming a planar sheet with narrow yolk stalks, basal myosin II instead accumulates into segmented patterns at cell-cell interfaces (Fig. 5A) (Thomas and Wieschaus 2004). This disrupted basal actomyosin organization makes *scra* RNAi embryos a useful system for exploring the role of basal tissue mechanical properties in directing morphogenetic behaviors. Thus, we predicted that furrowing direction bias and bending behavior would both be reverted by *scra* RNAi treatment.

**Figure 5.**
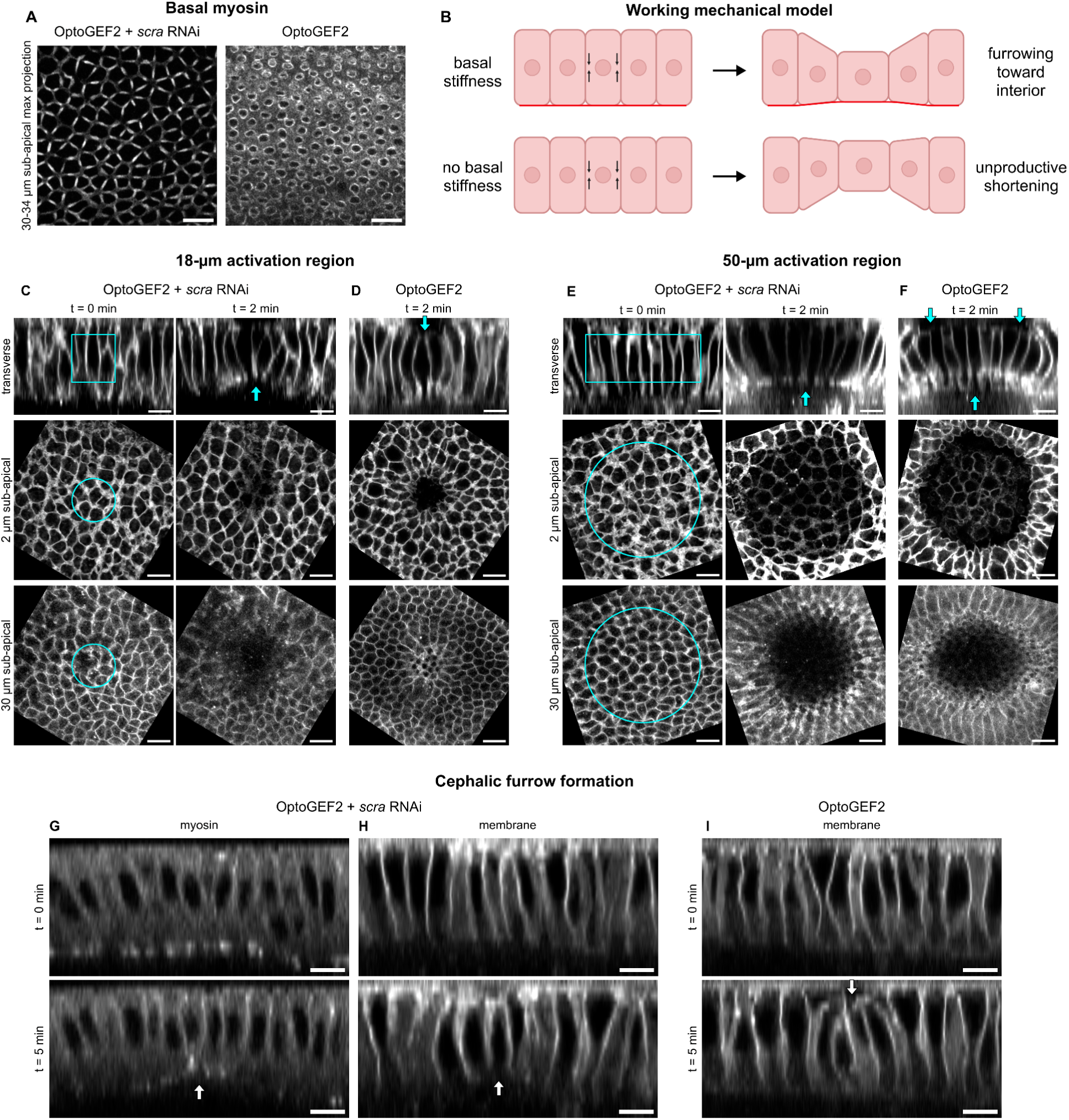
Perturbing basal myosin via *scraps/anillin* RNAi changes tissue furrowing and bending behaviors. **(A)** Basal myosin II at the onset of gastrulation in unactivated OptoGEF2 + *scra* RNAi embryos (left) and unactivated OptoGEF2 control embryos (right). **(B)** Schematic: We hypothesize that a stiff basal layer is required to properly orient cephalic furrow formation and to translate cell shortening into productive tissue furrowing. **(C, D)** Two-photon lateral activation of an 18-µm diameter region in embryos expressing OptoGEF2 + *scra* RNAi (C) or OptoGEF2 alone (D). In a *scra* RNAi background, cells shorten toward their apical surface (visualized by mCh-iLID-CaaX) and form an unproductive furrow. **(E, F)** Two-photon lateral activation of a 50-µm region in embryos expressing OptoGEF2 + *scra* RNAi (E) or OptoGEF2 alone (F). In a *scra* RNAi background, cells shorten without tissue-scale bending. **(G, H)** In unactivated embryos expressing OptoGEF2 + *scra* RNAi, myosin still accumulates in cephalic furrow initiator cells (G), but these cells now shorten toward their apical surfaces (H). n = 4/4 embryos show inverted CFF **(I)** In unactivated control embryos expressing OptoGEF2 alone, CFF occurs normally and cells shorten toward their basal surfaces. n = 0/3 embryos show inverted CFF. In C, D, E, F, H, I, mCh-iLID-Caax is the membrane marker. In A, G, myosin is visualized with sqh-mCh. Scale bars are 10 µm.

Consistent with our predictions, small lateral activation regions in OptoGEF2 + *scra* RNAi embryos caused shortening from the cell basal surface instead of from the apical surface, which was seen in embryos expressing OptoGEF2 alone (Figs. 5C, 5D). Large activation regions in OptoGEF2 + *scra* RNAi embryos caused shortening from the basal surface, but no overall tissue curvature, as demonstrated by the absence of the apical indentations that characterized bending in OptoGEF2 embryos (Figs. 5E, 5F). This suggests that a basal actomyosin layer is required for productive cell furrowing in response to lateral activation (Fig. 5B) and is the source of the bending phenotypes observed in large activation regions (Fig. 4G, left).

To understand how the disrupted basal layer in *scra* RNAi embryos affects normal lateral contraction-driven morphogenesis, we imaged cephalic furrow dynamics in OptoGEF2 + *scra* RNAi embryos without any activation. Notably, while lateral actomyosin still accumulated in cephalic furrow initiator cells (Fig. 5G), it now frequently caused these initiator cells to shorten from their basal surfaces rather than their apical surfaces (Figs. 5H, 5I). This was not a result of broadly disrupted epithelial morphogenesis, as ventral furrow formation still occurs normally in *scra* RNAi embryos (Goldner et al. 2023 *Preprint*), suggesting that the perturbed phenotype is particular to mechanisms driving cephalic furrow formation.

Overall, in both our optogenetic perturbations and endogenous cephalic furrow formation, productive lateral contraction-driven tissue furrowing towards the interior of the embryo requires a stiff basal layer. Although lateral contractility is the active force driving furrowing, passive basal stiffness seems to provide the symmetry-breaking mechanical cue that orients it. However, these basal mechanical properties have the additional effect of imposing a size constraint on lateral contraction-driven furrows, beyond which tissues bend instead of coherently furrowing.

## Discussion

Here, we developed two optogenetic tools based on the endogenous tagging of different RhoGEFs with a component of a light-activated heterodimer. To our knowledge, these are the first tools directly manipulating endogenous drivers of morphogenesis in *Drosophila*. They complement a growing body of optogenetic tools permitting light-directed control of endogenous protein activity (Redchuk et al., 2020, Manoilov et al., 2021, Kook et al., 2023).

### Advantages and limitations of tools manipulating endogenous RhoGEFs

Our work suggests that RhoGEFs are amenable to the presence of fluorescent and optogenetic protein tags without perturbing their function, consistent with a recent study (Di Pietro et al., 2023). The endogenous nature of these tools allowed us to accurately quantify our optogenetic input by accounting for expression variability of only the iLID component. In theory, a dose could be precisely quantified in any optogenetic system by measuring both component levels (Natwick and Collins 2021), but this requires two dedicated fluorescence channels and is prohibitively difficult for optogenetics applications. Both of the dose quantification strategies we present here can be applied to a variety of fluorescent imaging setups and activation methods.

The low endogenous GEF expression prevented us from imaging dynamics of GEF recruitment in response to 2P activation due to weak fluorescence signal and photobleaching. Despite this, we were able to validate our recruitment protocol via observation of 1P OptoGEF2/OptoCysts recruitment, 2P OptoSOS recruitment, the requirement for both tool components in eliciting a shortening response, and the failure of laser power alone to explain shortening behavior (Figs. S2 and S6). Moreover, in the experiments presented in the main text, we did not observe phenotypes indicative of IR-induced tissue damage (Fig. S2) (Nishizawa *et al*. 2023).

These tools are complementary to, but do not substitute for, transgenic OptoRhoGEFs based on GEF catalytic domains. Endogenous tools are well suited for studying endogenous full-length protein activities, manipulating morphogenesis in a healthy embryonic background, and quantitatively specifying cell and tissue shapes. Comparing the two tools developed here, OptoGEF2 drives larger deformations, while OptoCysts has improved localization of myosin recruitment. Nonetheless, both endogenous tools, particularly OptoGEF2, are limited by endogenous protein expression and localization patterns and potential off-target binding partners, making them less suitable for induction of apical constriction or for general, arbitrarily localized, on-target Rho pathway activation. Future iterations of these tools may benefit from a brighter fluorescent tag, a less diffusive membrane anchor (Natwick and Collins 2021), or improved two-photon activation efficiency, for example, by directing absorption through a FRET-coupled fluorophore (Kinjo et al., 2019).

### Regulatory logic of RhoGEF2

We observed that OptoGEF2, but not OptoCysts, produced unexpected myosin localization patterns - namely, subnuclear circumferentially constricted rings (Fig. 1E, upper middle) and persistent basal myosin (Fig. 1E, lower right). This resulted in transient lateral pinching (Fig. 1G) and an exacerbated bending phenotype (Figs. 4C and S6A). We note that this does not simply arise from basal light leakage or iLID diffusion, as activating a 5-10 µm sub-apical volume produced the same phenotypes as activating a 5-25 µm sub-apical volume (Figs. S7E, S7F), and these length scales are inconsistent with those expected from two-photon axial spreading (Dong et al., 2003) or from iLID-CaaX diffusion (Natwick and Collins 2021). Moreover, the absence of these behaviors after OptoCysts or RhoGEF2(DHPH)-Cry2 activation (Izquierdo et al., 2018, Popkova et al., 2024) suggests that they reflect a specific feature of endogenous full-length RhoGEF2.

Our optogenetic manipulations of endogenous RhoGEF2 localization complement traditional biochemistry approaches in dissecting RhoGEF2 regulatory logic. In the *Drosophila* primary epithelium, RhoGEF2 is sequestered at MT plus ends in order to prevent its constitutive activity (Rogers et al., 2004, Lin et al., 2022). Selective employment of RhoGEF2 in different developmental contexts frequently requires both its release from MTs and its anchoring to a new subcellular membrane compartment. During ventral furrow formation, RhoGEF2 is relieved from MT sequestration by Gα_12/13_/Concertina (Cta) binding, then anchored at apical cell membranes by T48 (Parks and Wieschaus 1991, Rogers et al., 2004, Kölsch et al., 2007). During *Drosophila* primordial germ cell (PGC) migration, RhoGEF2 is released from MTs by AMPK phosphorylation of a MT-binding domain, then anchored to polarized PGC rears to drive oriented amoeboid migration (Lin et al., 2022).

Surprisingly, we find that explicit release of RhoGEF2 from MTs is not required for its recruitment or for rapid induction of large cell shape changes. Providing membrane anchoring sites is sufficient to promote RhoGEF2 relocalization and contractile activity. The relatively weak apical constriction response generated by apical OptoGEF2 activation is reminiscent of *Cta* mutants, which still show a residual level of apical constriction and furrowing due to T48 binding (Parks and Wieschaus 1991, Kölsch et al., 2007). In this context, active iLID plays a similar role to T48 - both redirect a non-sequestered subpopulation of RhoGEF2 to the apical surface, where they drive a small amount of apical constriction that is insufficient for robust tissue invagination. It is unknown what interactions induce RhoGEF2-dependent lateral contraction in the cephalic furrow, but our work suggests that lateral membrane anchors are sufficient to drive cell shortening during cephalic furrow formation.

RhoGEF2 PH domain-mediated positive feedback with active Rho1-GTP potentiates RhoGEF2 recruitment and activity (Chen et al., 2010, Medina et al., 2013, Rich et al., 2020, Lin et al., 2022). Given that Cysts also acts via Rho1, it’s possible that Cysts recruitment to membranes results in additional RhoGEF2 recruitment and strengthens the contractile response. However, based on the qualitative differences between OptoGEF2 and OptoCysts activation described above, it seems that RhoGEF2 recruitment is not the dominant mechanism of OptoCysts-driven contraction.

We propose the following account of OptoGEF2-specific behaviors. Upon iLID activation, RhoGEF2 is recruited along lateral membranes, especially at perinuclear positions via enhanced capture of punctate RhoGEF2 from membrane-apposed MT bundles surrounding cell nuclei (Warn and Warn 1986, Fig. S4B). This drives subnuclear circumferential constriction, possibly in combination with additional RhoGEF2-specific interactions or cytoskeletal structural elements particular to subnuclear positions (Krueger et al., 2019b). Once liberated, RhoGEF2 diffuses laterally and searches for a new anchoring site, where it may encounter Slam, a RhoGEF2 anchor responsible for its localization at the cellularization front (Wenzl et al., 2010). Slam traps MT-liberated RhoGEF2 basally, resulting in basal myosin accumulation and additional basal tension. Combined with further genetic and biochemical manipulations, endogenous tools will be useful for testing these ideas and understanding other aspects of endogenous RhoGEF behavior.

### Design principles of epithelial furrowing

Through our quantitative studies, we uncovered several key principles of lateral contraction-driven tissue morphogenesis. We found that furrows formed by this mechanism can have their morphologies specified by grading levels of RhoGEF membrane anchors, which acts by modulating the amount of membrane-bound RhoGEF. However, patterning of tissue apical-basal mechanical properties plays a crucial role in determining the resulting tissue deformations. A stiff basal actomyosin layer, present at the onset of gastrulation, not only provides a symmetry-breaking cue to orient tissue furrowing, but also constrains the size of productive furrows.

Although residual basal myosin in OptoGEF2 embryos may exacerbate these behaviors, they were also observed in OptoCysts embryos and were dependent on an intact basal actomyosin network. This stiff basal network is characteristic of cellularization (Cheikh et al., 2023), but is a transient state in gastrulation and persists only for the first several minutes of ventral and cephalic furrow formation. Nonetheless, apical-basal asymmetry is a defining property of epithelia (Buckley and St Johnston 2022), and the mechanics described here are potentially representative of many epithelia with mature basement membranes. In other contexts, a stiff apical surface could direct furrowing behaviors, or buckling and wrinkling processes may emerge in the absence of an explicit symmetry-breaking cue.

The ventral and cephalic furrows employ similar strategies to permit cell internalization past a stiff basal surface. Both the presumptive mesoderm and cephalic furrow initiator cells undergo precocious loss of basal actomyosin in order to couple cellular constriction to basal expansion and invagination (Dawes-Hoang et al., 2005, Polyakov et al., 2014, Spencer et al., 2015, Popkova et al., 2024). Ectopic local stiffening of the basal tissue surface via optogenetics prevents both ventral and cephalic furrow internalization (Krueger et al., 2018, Popkova et al., 2024).

Our results suggest that preceding relaxation, cephalic furrow formation takes advantage of the stiff tissue basal surface to orient cell shortening toward the embryo interior. This may prime cells to invaginate once basal myosin is lost, either by stabilizing the shortened cells via plastic deformation (Molnar and Labouesse 2021) or by overcoming an energy barrier associated with cell movement (Bi et al., 2014, Takeda et al., 2020). Alternatively, given that basal relaxation is sufficient to cause cell ingression in the primary epithelium (Popkova et al., 2024), that cell pressure differentials play major roles in determining cephalic furrow cell shapes (Niloy et al., 2023), and that hydrostatic pressure excess can enhance tissue folding past a compliant surface (Guerra Santillán et al. 2024b *Preprint*), the role of the basal sheet may instead be to increase hydrostatic pressure in response to cell contraction.

Surprisingly, we find that lateral contraction does not scale well as a furrowing mechanism in the primary epithelium. When a large group of cells shortens at once, restricted basal area expansion causes tissues to adopt curved shapes, forcing central cells to move toward the vitelline membrane instead of toward the embryonic interior. Although reminiscent of buckling processes in other epithelia and developing systems (Savin et al., 2011, Nelson 2016, Trushko et al., 2020, Wang et al., 2024), this process is distinct because it requires no external forces and exhibits no sudden transition from a non-buckled to a buckled state. This continuous change in curvature as a function of lateral contraction has been described theoretically (Guerra Santillán et al., 2024a), but to our knowledge, has not been demonstrated experimentally. It will be interesting to re-examine wedged cell shapes and curved epithelial morphologies in embryos to see if this curvature mechanism is at play during development.

Endogenous optogenetic tools, combined with the dose quantification strategies we present here, provide a powerful toolbox to interrogate epithelial morphogenesis. We build upon the spatial and temporal control inherent to optogenetic systems by additionally permitting quantitative induction of tool activation and biomechanical responses. Our work lays a foundation for the precise modulation of contractile forces from the subcellular to tissue scales, allowing for careful interrogation of developmental mechanics and control of 3D shapes of engineered living systems and materials.

## Acknowledgements

We thank Stas Shvartsman and Liz Gavis for their contributions to conceptualization of the endogenous optogenetic tools and for helpful discussions. We thank Bex Pendrak, Sameer Thukral, and Kasza Lab members for helpful discussions. Stocks obtained from the Bloomington Drosophila Stock Center (NIH P40OD018537) were used in this study. This work was performed in part at the Live Imaging and Bioenergetics Facility at the Advanced Science Research Center at The Graduate Center of the City University of New York. This work was supported by NIH Grant R35GM138380 to KEK; KEK holds an NSF CAREER Award, Packard Fellowship, and Sloan Research Fellowship in Physics.

## Methods

### Cloning

To make pBI-UASp-mCherry-iLID-CaaX, pBI-UASp-Venus-iLID-CaaX, and pBI-UASp-iLID-Caax, Venus and iLID-CaaX were amplified from pLL7.0: Venus-iLID-CAAX (Addgene #60411, gift from Brian Kuhlman) and mCherry was amplified from pBI-UASp-mCherry-Cry2phr-RhoGAP71E (Herrera-Perez et al., 2021). Fragments were cloned into a pENTR/D-Topo vector (Life Technologies, Carlsbad, CA) using the NEBuilder HiFi DNA Assembly Kit (New England BioLabs, Ipswich, MA) and recombined into a UASp-attB destination vector (gift of F. Wirtz-Peitz) using the Gateway cloning system (Life Technologies, Carlsbad, CA).

To make pHD-RhoGEF2-tdTomato-Cry2-DsRed, pHD-RhoGEF2-tgRFPt-SspB-DsRed, pHD-RhoGEF2-SspB-DsRed, pHD-SspB-tgRFPt-DsRed-Cysts, and pHD-SspB-DsRed-Cysts, SspB and tgRFPt were amplified from pLL7.0: tgRFPt-SSPB WT (Addgene #60415, gift from Brian Kuhlman), tdTomato was amplified from pCAG-tdTomato (Addgene #83029, gift from Angelique Bordey), Cry2 was amplified from pBI-UASp-mCherry-Cry2phr-RhoGAP71E (Herrera-Perez et al., 2021), DsRed was amplified from pScarlessHD-sfGFP-DsRed (Addgene #80811, gift from Kate O’Connor-Giles), and homology arms were amplified from genomic DNA of vas-Cas9 III (BDSC #51324). We used FlyBase (release FB2020_06) to obtain RhoGEF2 and Cysts homology sequences. Fragments were cloned into a pScarlessHD-sfGFP-DsRed donor vector (Addgene #80811, gift from Kate O’Connor-Giles) using the NEBuilder HiFi DNA Assembly Kit (New England BioLabs, Ipswich, MA).

Donor plasmid PAM sites were mutated using the QuikChange II XL Site-Directed Mutagenesis Kit (Agilent Technologies, Santa Clara, CA). The gRNA-encoding plasmids pU6-BbsI-chiRNA-RhoGEF2 and pU6-BbsI-chiRNA-Cysts were made by insertion of gRNA oligos into pU6-BbsI-chiRNA (Addgene #45946, gift from Melissa Harrison and Kate O’Connor-Giles and Jill Wildonger) using the Q5 Site-Directed Mutagenesis Kit (New England BioLabs, Ipswich, MA).

### Fly stocks

*Drosophila melanogaster* was cultured at 23 °C using standard techniques. Crosses were maintained in the dark at 23 °C. Imaging was done in the dark at room temperature (∼21 °C). Crosses and embryos were handled in dark rooms with all light sources covered by a Red 25 Wratten Filter (Kodak, Rochester, NY).

iLID constructs under a UASp promoter were expressed with a maternal α-tubulin matα-tub15 Gal4-VP16 driver (mat15, gift from D. St Johnston). CRISPR/Cas9 knock-ins at RhoGEF2 and Cysts loci were done following the flyCRISPR method (https://flycrispr.org/) (Gratz et al., 2014). Guide RNAs were designed using the flyCRISPR target finder (http://targetfinder.flycrispr.neuro.brown.edu/). Homology donor and gRNA plasmids were co-injected into vas-Cas9 III (BDSC #51324) (BestGene Inc, Chino Hills, CA). After homology-mediated cassette insertion, the 3xP3-DsRed selectable marker was flipped out by crossing to nos-PBac (III) (Keenan et al., 2020). Transgenic iLID lines were generated by PhiC31-directed integration into an attP2 docking site (BDSC #8622) (BestGene Inc, Chino Hills, CA).

The fly stocks generated for this study were RhoGEF2-tgRFPt-SspB, RhoGEF2-SspB, RhoGEF2-tdTomato-Cry2, SspB-tRFP-Cysts, SspB-Cysts, UASp>iLID-CaaX (III), UASp>Venus-iLID-CaaX (III), UASp>mCherry-iLID-CaaX (III). Other stocks used in this study were mat15 (III), sqh>sqh-mCherry (III) (BDSC #59024, donated by Beth Stronach), UASp>OptoSOS (II) (BDSC #95111, donated by Jared Toettcher), UASp>*scra*(RNAi) (III) (BDSC #41841, Transgenic RNAi Project), sqh>gap43-mCherry (III) (gift from Adam Martin, Martin et al., 2010).

### Live imaging and photomanipulation

Embryos were collected in apple agar plates in the dark, dechorionated for 2 min in 50% bleach, washed with water for 1 min, selected under Halocarbon oil 27 (Sigma-Aldrich, St. Louis, MO), and mounted in a 50/50 mixture of Halocarbon oil 27 and Halocarbon oil 700 (Sigma-Aldrich, St. Louis, MO) between a glass coverslip (VWR International, Radnor, PA) and an oxygen-permeable membrane (YSI, Yellow Springs, OH). For most experiments, embryos were imaged and activated on a Zeiss LSM 880 NLO upright two photon confocal microscope, equipped with a diode laser for 561 nm excitation, an argon laser for 488 nm excitation, a Mai Tai DeepSee 2 photon laser for 690-1040 nm excitation, an Airyscan detector with a FAST module, and a Plan-Apochromat 40x/1.0 NA water objective. For cephalic furrow imaging (Fig. 5G-5I) and RhoGEF2 Airyscan imaging (Fig. S4), embryos were imaged on a Zeiss LSM 880 inverted microscope, equipped with a diode laser for 561 nm excitation, an Airyscan detector with a FAST module, and a C-Apochromat 40x/1.2 NA water objective (for CF imaging) or a Plan-Apochromat 63x/1.40 NA oil objective (for RhoGEF2 imaging).

Embryos were activated on their dorsal surfaces, at the midpoint between the anterior and posterior poles. Embryos were activated at the onset of gastrulation, when cells were between 32 and 35 µm long. During lateral activation experiments, embryos were activated by scanning a 927-nm Mai Tai laser in a circular region through a cylindrical stack extending from 5-25 µm below the apical cell surface, sampled every 1 µm, with a pixel dwell of 1.85 µs, a pixel spacing of 0.12 µm, and a laser power between 0-70% (0-75 mW at sample plane), for 5 total iterations. During apical activation experiments, a 927-nm Mai Tai laser was scanned in a circular region at a single apical slice, with a pixel dwell of 1.85 µs, a pixel spacing of 0.12 µm, a laser power of 45% (47 mW at sample plane), for 3 iterations per frame, repeated every frame (every 22 s). Movies tracking membrane markers and myosin were taken as 44-µm stacks with 2 µm slice spacing, taking 22 s/frame. Vitelline membrane micro-perforation was done by scanning a 840-nm laser at 100% power (nominally 2600 mW at the laser source) over a 5 µm x 10 µm patch of the vitelline membrane, sampled every 0.12 µm in xy and every 0.5 µm in z, for 1 iteration (Popkova et al., 2024).

### Furrow segmentation and tracking

Microscopy data were viewed in ImageJ (Schneider et al., 2012). Optogenetically induced furrows were quantified by segmenting tissue apical and basal boundaries. Tissue apical surfaces were segmented using an ilastik autocontext workflow trained on manually annotated data with all color/intensity, edge, and texture features selected (Berg et al., 2019). Output pixel probabilities were fed into an ilastik object classification workflow with 2x2 smoothing, hysteresis thresholding (core probability 0.5, edge probability 0.35), and were classified using the following features: convexity, number of defects, convex hull center, size in pixels, mean intensity, and histogram of intensity in neighborhood (10 x 10). Segmentations and classifications were manually confirmed and corrected when necessary. Due to poor basal membrane signal, basal tissue boundaries were segmented manually in ImageJ. Tissue movement was tracked and corrected for using PIVlab (FFT window deformation, 128px first pass, 64px second pass, 32px third pass) (Thielicke and Stamhuis 2014).

### Furrow quantification

Using the segmented tissue boundaries, cell height at each point in the tissue was calculated as the difference between the z-position of the apical tissue boundary and that of the basal tissue boundary. To avoid artifacts from embryo curvature, this was only done at coverslip-adjacent flattened portions of the embryo. A reference cell height curve, representing what cell height would be in the absence of activation, was constructed by using non-activated portions of the embryo to calculate an embryo-specific cellularization rate, which was used to estimate expected cell lengthening through the end of the experiment. The shortening at each point of the tissue was then calculated as the difference between the instantaneous cell height and the reference cell height curve. “Average shortening” is measured as the largest cell shortening at any time point in the experiment, averaged across all points within the activation region.“Maximum shortening” is measured as the largest cell shortening at any time point in the experiment, measured at any point within the activation region. The movement of the activation region was tracked and accounted for via PIV, as described above. Furrowing and recovery timescales were calculated by finding the time point at which the maximum cell height reached 1/e =37% of its final shortened or recovered height, interpolating between adjacent time points when necessary.

### Activation quantification

iLID activation levels were quantified via the fluorescence signal emitted during the first activation time point (20-µm stack, 5-25 µm below the embryo apical surface, sampled every µm, 21 slices total, shown in Fig. 2A’). A standard fluorescence curve was calculated as the average fluorescence intensity at each slice, averaged across all experiments. For each experiment, the normalized fluorescence at each slice was calculated as that experiment’s fluorescence at that slice divided by the standard curve’s fluorescence at that slice. The final normalized activation level for each experiment was calculated as the average of the experiment’s normalized fluorescence across all slices. This was used as the activation magnitude for the experiments in the main text. For the alternative activation magnitude calculation (Fig. S6), the same procedure was instead applied to the final pre-activation time point (Fig. 2A) to get an iLID expression level. This expression level was converted into a proxy for activation magnitude by multiplying it with the square of the laser power, calculated as the product of the user-input laser power (% max) with the maximum laser power (mW) measured at the sample plane.

### Viability studies

Cages were kept at 23 °C in either 24-h dark or 24-h light. Embryos were collected into apple agar plates over a 72-hours period, split into an 8-hour and a 16-hour collection block every day. After collection, embryos were arranged in a grid for counting, and plates were laid out to be exposed to either 24-h dark or 24-h light. 24-48 hours later, the number of hatched embryos was counted.

### Endogenous expression quantification

Embryos homozygous for RhoGEF2-tgRFPt-SspB, SspB-tgRFPt-Cysts, and vas-Cas9 (III) were mounted as described. Using bright-field transmitted light, embryos were selected that were in late stage 5/early stage 6 and had their dorsal surfaces facing the coverslip. A single fluorescent image was taken 5 µm below the vitelline membrane over a 15-s scan period using the 561-nm laser line. A fluorescence signal was calculated by measuring the average fluorescence intensity across the tissue, leaving a 5-µm buffer from the vitelline membrane to exclude vitelline membrane autofluorescence. A background fluorescence level was calculated as the average fluorescence across 5 measured vas-Cas9 (III) embryos. The expression level of each RhoGEF2-tgRFPt-SspB and SspB-tgRFPt-Cysts embryo was calculated as the average fluorescence of that embryo minus the background vas-Cas9 (III) fluorescence value.

### Figure preparation

All figures were prepared in Figma. Graphs were made in Python using Matplotlib (Hunter 2007).

### Statistical analysis

Statistical analyses were done in Python using SciPy (Virtanen et al., 2020). ANOVAs and t-tests were performed as noted. Tests for coincidence of two lines were done as described in Glantz 2012.

