## Supplementary Material for "Endogenous OptoRhoGEFs reveal biophysical principles of epithelial tissue furrowing"

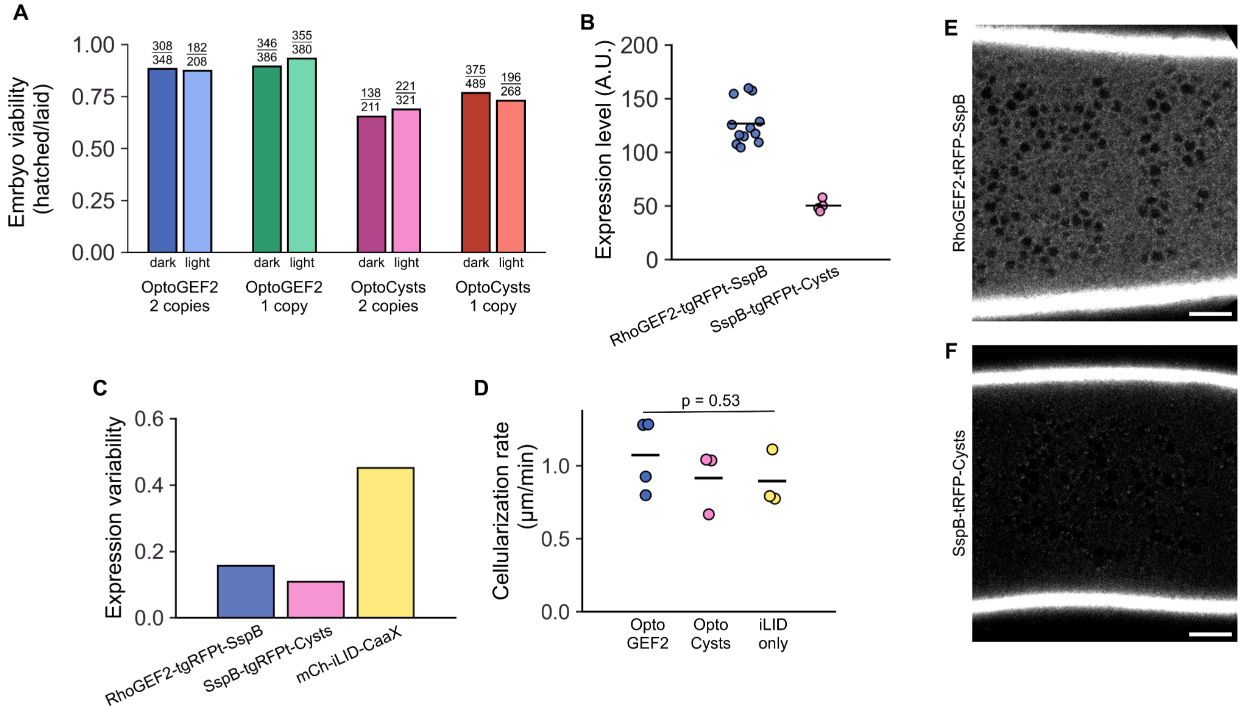


**Figure S1**. Flies expressing endogenous OptoRhoGEFs are viable, healthy and show little variability. **(A)** Eggs laid by mothers expressing mCh-iLID-CaaX and either 1 or 2 copies of RhoGEF2-tgRFPt-SspB or SspB-tgRFPt-Cysts are viable when incubated in either dark or light. **(B)** Expression levels of RhoGEF2-tgRFPt-SspB and SspB-tgRFPt-Cysts in the early stage 6 dorsal epithelium, as measured by tgRFPt fluorescence. Expression levels are different at $p < 10^{-5}$ (t-test). **(C)** Endogenous GEF expression shows less variability than UAS-driven mCh-iLID-Caax expression, as quantified by the standard deviation of the expression level divided by mean expression level. **(D)** Cell lengthening rates during the fast stage of cellularization are not different between embryos expressing OptoGEF2, OptoCysts, or mCh-iLID-CaaX only ($p = 0.53$, ANOVA). **(E, F)** Sample images of RhoGEF2-tgRFPt-SspB (E) or SspB-tgRFPt-Cysts (F) used to calculate expression levels in (B), presented using the same display settings. Scale bars are 20 μm.

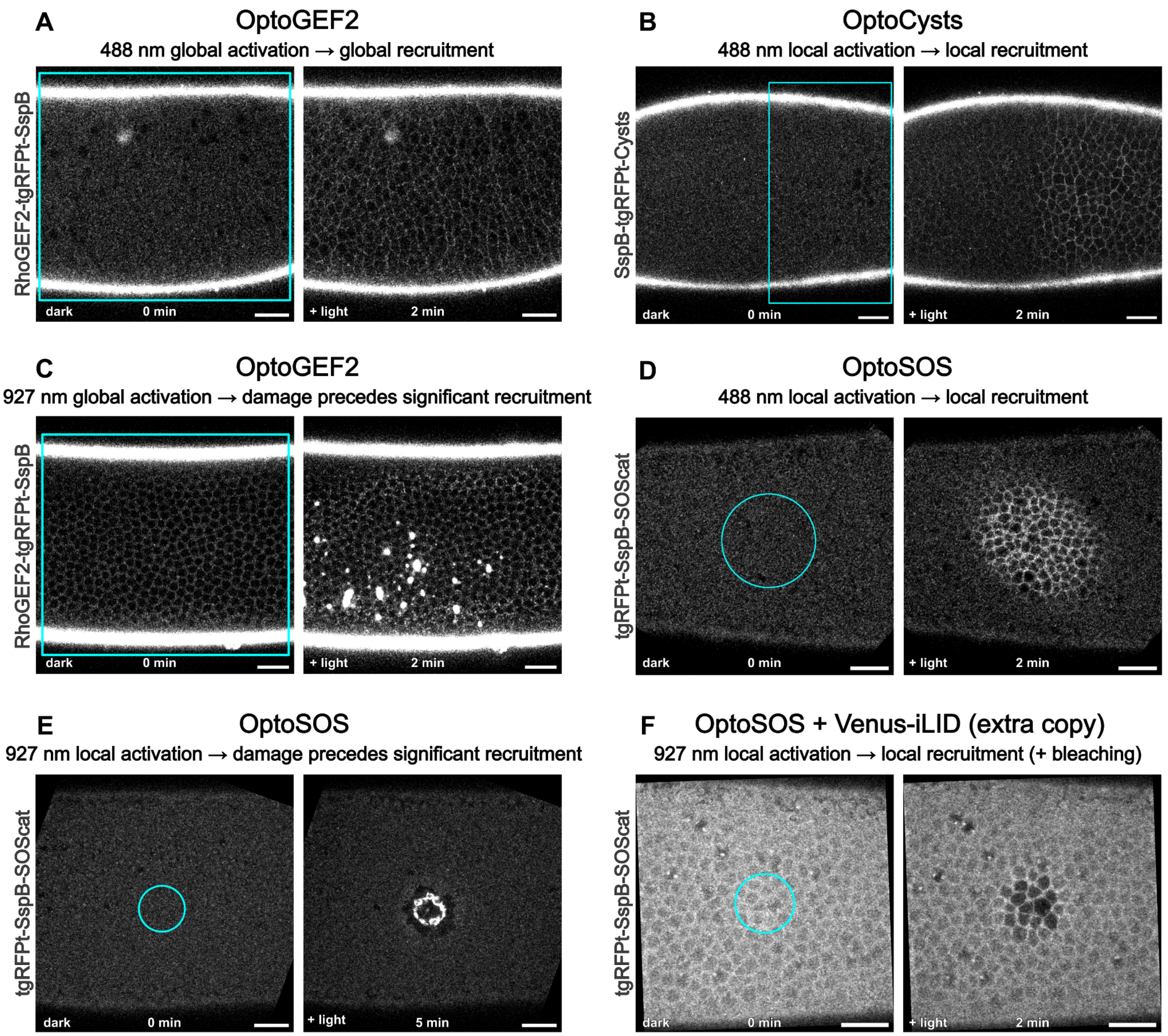


**Figure S2**. Expressing iLID in excess of SspB permits strong recruitment with weak activation protocols. **(A)** In embryos co-expressing RhoGEF2-tgRFPt-SspB and iLID-CaaX, continuous global 488-nm illumination causes global RhoGEF2 recruitment to cell membranes. **(B)** In embryos co-expressing SspB-tgRFPt-Cysts and iLID-CaaX, continuous local 488-nm illumination causes local Cysts recruitment to cell membranes within the illuminated area. **(C)** In embryos co-expressing RhoGEF2-tgRFPt-SspB and iLID-CaaX, global 927-nm illumination (continuous scanning, 81 mW) causes tissue ablation before any recruitment becomes visible. **(D)** In embryos co-expressing tgRFPt-SspB-SOScat and iLID-CaaX, continuous local 488-nm illumination causes local SOScat recruitment to cell membranes within the illuminated area. **(E)** In embryos co-expressing tgRFPt-SspB-SOScat and iLID-CaaX, 927-nm local activation (every 15 s for 4 s, 79 mW) causes tissue ablation after 5 minutes before any recruitment becomes visible. **(F)** In embryos co-expressing tgRFPt-SspB-SOScat, iLID-CaaX, and an additional copy of Venus-iLID-CaaX, 927-nm local activation (every 7.5 s for 1 s, 76 mW) recruits SOScat to cell membranes within the illuminated area while causing additional bleaching, but without damaging the tissue. All slices are 6 µm below the apical surface. Scale bars are 20 µm.

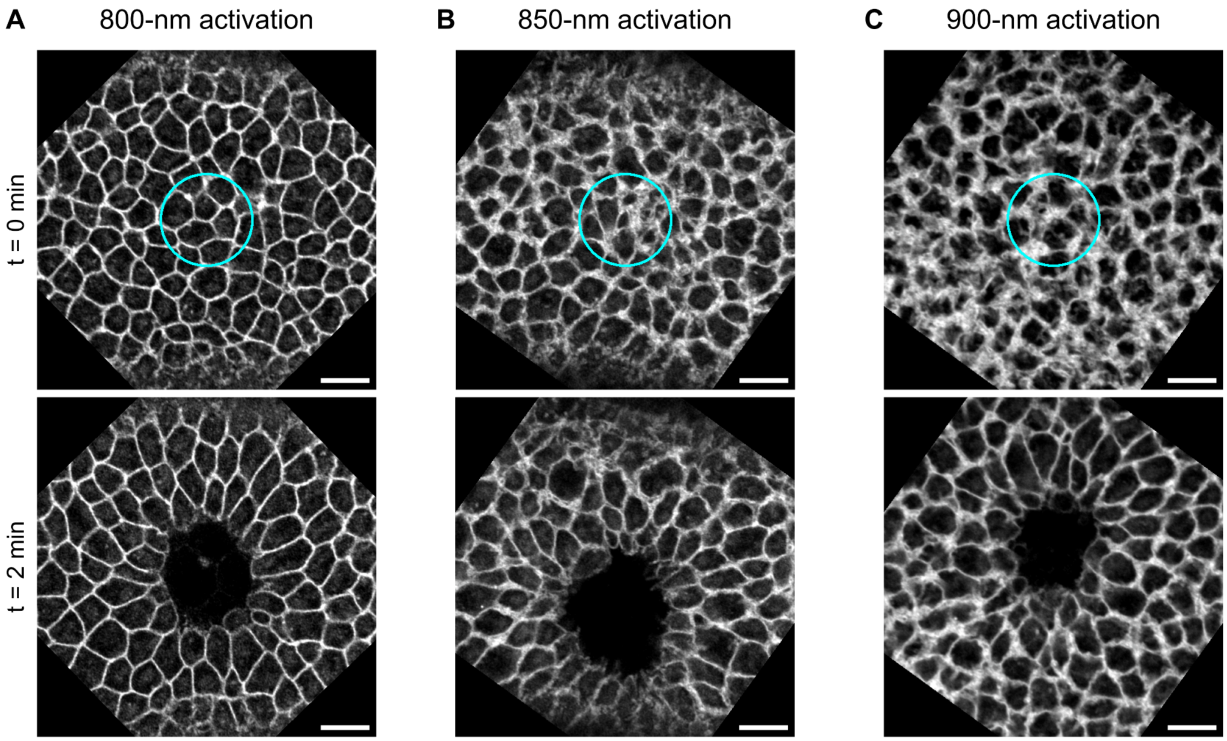


**Figure S3**. Cell shortening can be induced by a broad spectrum of IR illumination to activate OptoGEF2. **(A, B, C)** Embryo response to lateral RhoGEF2 recruitment using IR light of 800 (A), 850 (B), or 900 (C) nm wavelength. Membrane marker is mCh-iLID-CaaX. Slices are 2 μm below the apical surface. Scale bars are 10 μm.

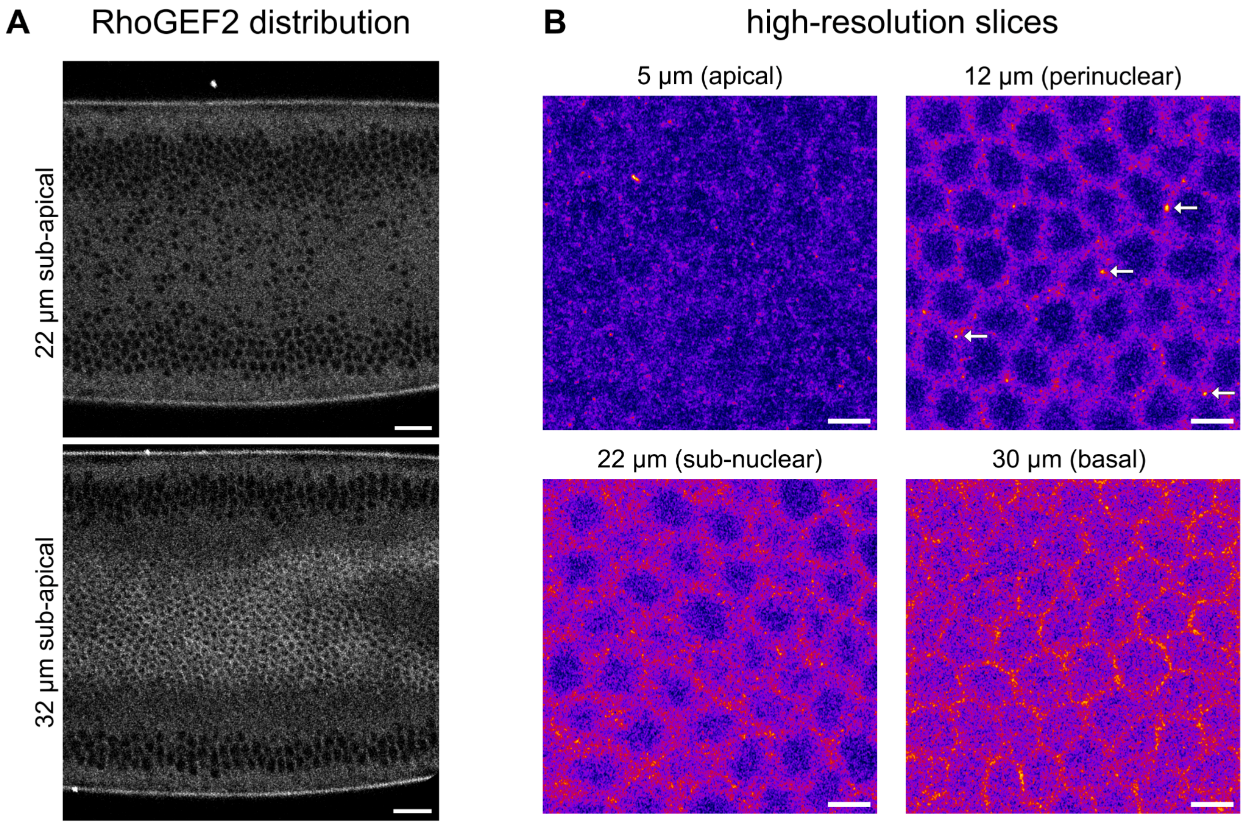


**Figure S4**. Endogenous RhoGEF2 is enriched at perinuclear puncta. **(A)** Cross-sections through the cellularizing dorsal primary epithelium of RhoGEF2-tdTomato-Cry2 embryos, taken when cells were 32 µm long. RhoGEF2 is enriched at the cellularization front (bottom), but otherwise displays a uniform cytoplasmic distribution (top). Scale bars are 20 µm. **(B)** Airyscan images of RhoGEF2 localization at various tissue depths, taken in adjacent portions in a single RhoGEF2-tdTomato-Cry2 embryo where the cellularization front had reached a depth of 32 µm. Apical cell regions contain small, sparse RhoGEF2 puncta. Perinuclear regions contain strong cytoplasmic RhoGEF2 puncta (marked with white arrows). Basal regions contain membrane-localized puncta. Scale bars are 5 µm.

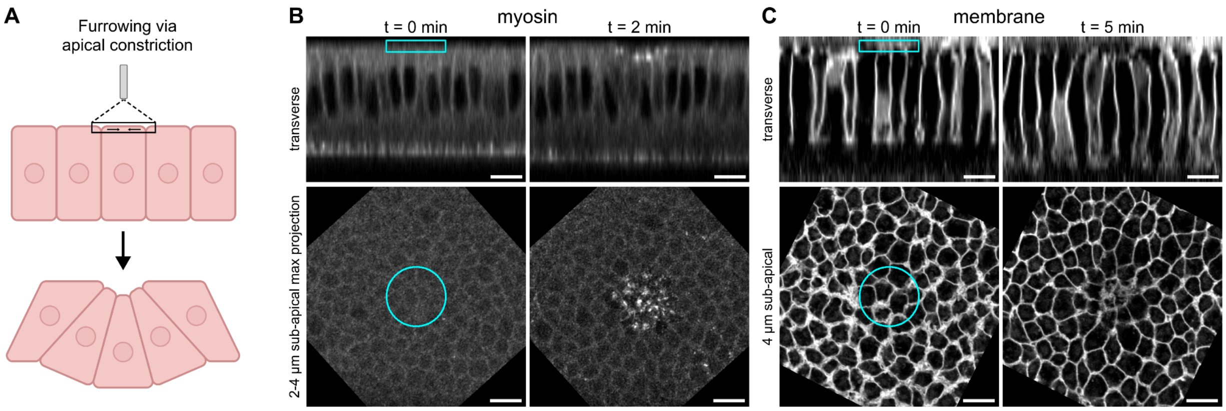


**Figure S5**. Apical OptoGEF2 activation leads to apical myosin accumulation and apical constriction but does not form a stereotypical furrow. **(A)** Schematic: Based on previous optogenetic tools and knowledge of endogenous furrowing processes, we expect that apical activation will form a furrow via apical constriction and basal expansion and ingression (Martin and Goldstein 2014). **(B)** In embryos co-expressing RhoGEF2-tgRFPt-SspB, Venus-iLID-CaaX, and sqh-mCherry, continuous apical 927-nm illumination leads to localized myosin recruitment to apical caps and formation of a small dimple on the apical surface. **(C)** In embryos co-expressing RhoGEF2-tgRFPt-SspB and mCherry-iLID-CaaX, continuous apical 927-nm illumination leads to the formation of a small dimple on the apical surface, accompanied by lateral, but not basal, expansion. Scale bars are 10 µm.

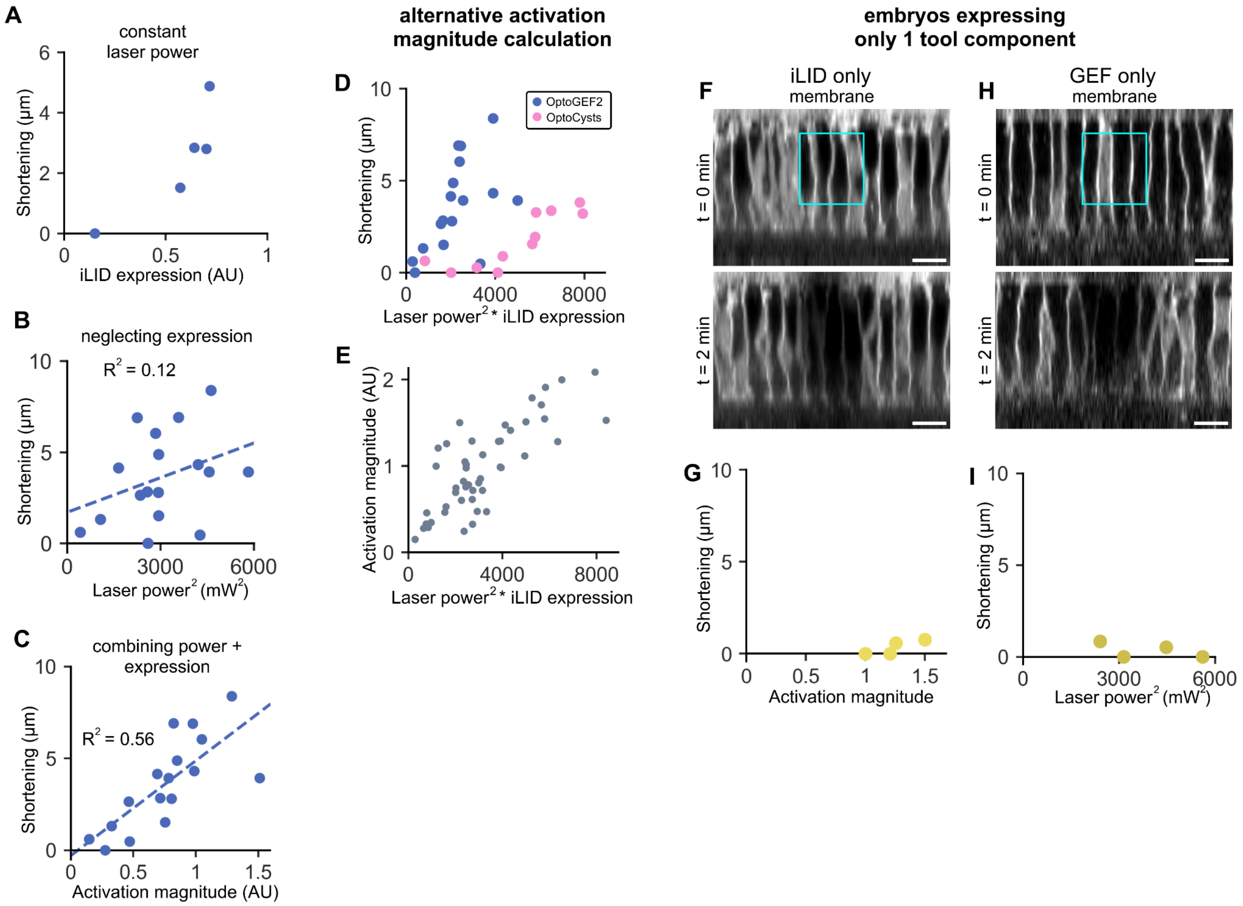


**Figure S6**. Shortening response depends on both activating laser power and iLID expression level. **(A)** At constant input laser power, cell shortening response depends on the level of iLID expression. **(B, C)** Cell shortening response is poorly predicted by activating laser power alone (B), but well predicted by an activation magnitude parameter (C), which is calculated by measuring emitted iLID fluorescence signal during the 927-nm activation step (Figure 2A') and combines the effects of activating laser power and iLID expression. **(D)** An alternative measure of activation magnitude, which also predicts cell shortening, can be calculated by separately measuring laser power at the 927-nm laser source and iLID expression in the 561-nm channel, and taking the product of the squared laser power with the level of iLID expression. **(E)** This alternative activation magnitude calculation, which is amenable to a wide range of activation setups, scales linearly with our standard activation magnitude parameter. **(F, G)** Embryos expressing only mCh-iLID-CaaX show no response to 927-nm lateral activation except for local photobleaching. **(H, I)** Embryos expressing only RhoGEF2-tgRFPt-SspB and a gap43-mCh membrane marker show no response to 927-nm lateral activation except for local photobleaching. Scale bars are 10 μm.

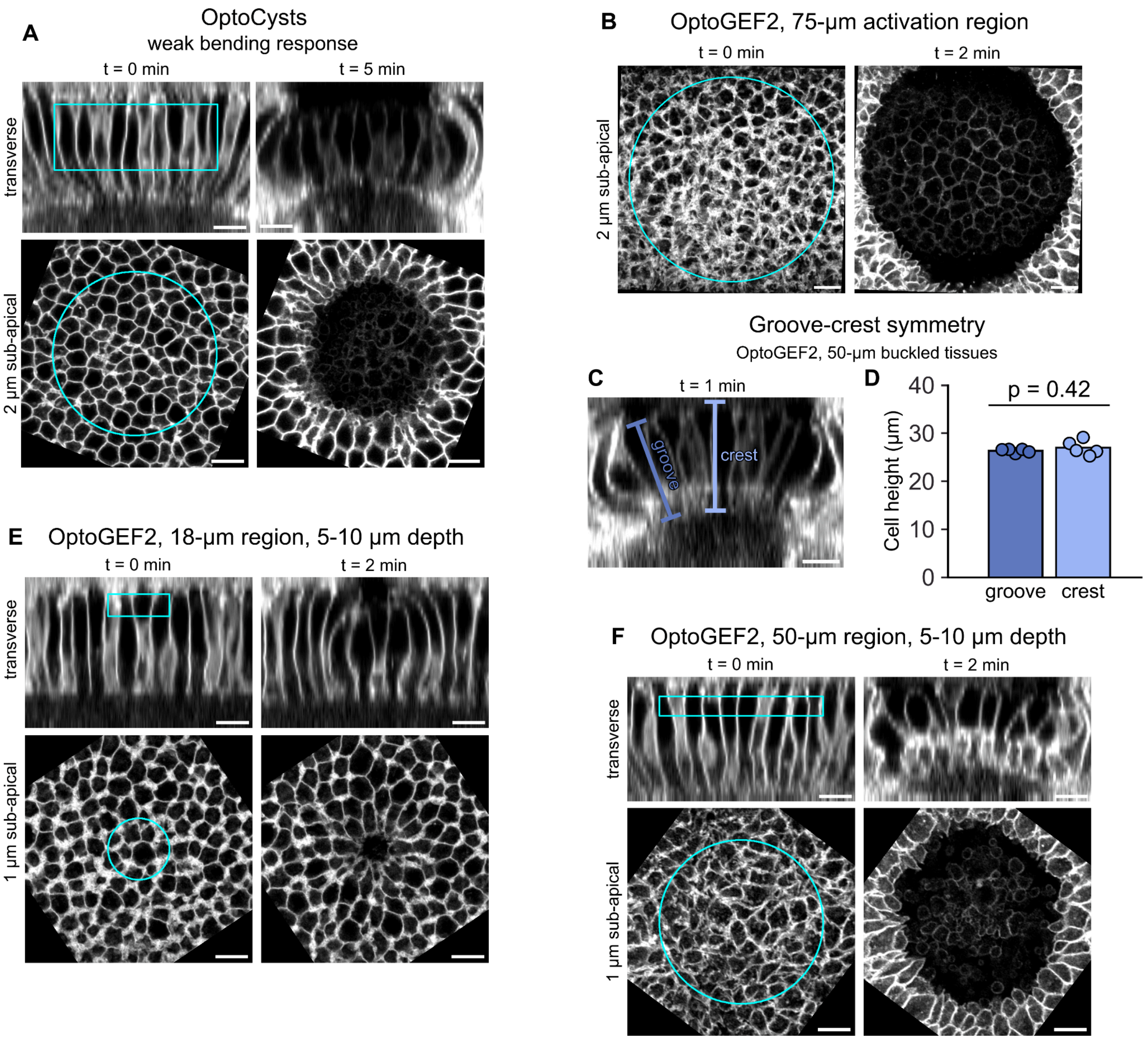


**Figure S7**. Tissue curvature is independent of the specific RhoGEF activated and is not the result of a wrinkling instability. **(A)** Lateral 927-nm activation of a 37-µm diameter cylindrical region in OptoCysts embryos results in a weak curvature response, qualitatively similar to OptoGEF2 embryos. **(B)** Lateral 927-nm activation of a 50-µm diameter cylindrical region in OptoGEF2 embryos produces a curved tissue with only a first-harmonic bending response, indicating that bending behavior is not periodic and is therefore not the result of a wrinkling instability. **(C)** In OptoGEF2 embryos that showed tissue bending after lateral activation of a 50-µm diameter cylindrical region, groove and crest heights were measured at 1 minute post-activation. **(D)** Cell heights in bent tissues are not significantly different at the groove and at the crest, suggesting that bending does not result from a wrinkling instability ($p = 0.42$, paired t-test). **(E)** In OptoGEF2 embryos, lateral activation of a 18-µm circle 5-10 µm below the apical surface produces similar bending behavior as lateral activation 5-25 µm below the apical surface (c.f. main text, Fig. 4). **(F)** In OptoGEF2 embryos, lateral activation of an 50-µm circle 5-10 µm below the apical surface produces similar bending behavior as lateral activation 5-25 µm below the apical surface (c.f. main text, Fig. 2). All images use a fluorescent mCh-iLID-CaaX membrane marker. Scale bars are 10 µm.
